# Using somatic data to aid germline clinical variant interpretation in developmental disorders

**DOI:** 10.64898/2026.06.17.732808

**Authors:** Katrina A Andrews, Matthew DC Neville, Iñigo Martincorena, Raheleh Rahbari, Helen Firth, Sarah Lindsay, Marc Tischkowitz, Matthew E Hurles

**Affiliations:** Wellcome Sanger Institute, Hinxton, Cambridge, CB10 1SA, UK; Department of Genomic Medicine, University of Cambridge, Cambridge Biomedical Campus, Cambridge, CB2 0QQ, UK

## Abstract

Accurate interpretation of rare germline variants remains a major challenge in developmental disorders (DD). Somatic mutation data represent a largely untapped source of evidence for germline variant classification. Identical or nearby mutations that drive positive selection when present in somatic tissues can cause developmental disorders when present in the germline. We integrated somatic mutation data from the Catalogue Of Somatic Mutations In Cancer (COSMIC), and healthy tissues (sperm and buccal epithelium) with germline variant datasets from ClinVar and large studies of de novo mutations in DD patients. Across 970 dominant DD genes, 195 have evidence of somatic selection, with a majority demonstrating concordant mechanisms between germline and somatic contexts. We benchmark the ability of somatic data to discriminate pathogenic from benign germline missense variation across dominant DD genes, identifying 145 genes in which somatic data are informative. The strongest utility is in altered-function genes where germline and somatic mechanisms are concordant, for example the RASopathy genes. In these genes, codon-level aggregation of somatic missense counts yields predictive performance comparable to computational predictors or MAVE assays (AUC-ROC 0.895 for somatic data, versus 0.893 for REVEL). Combining somatic features with computational scores improves discrimination further. Using likelihood ratios, we map COSMIC missense codon count thresholds onto American College of Medical Genetics and Genomics/Association for Molecular Pathology (ACMG/AMP)-style evidence strengths, showing that somatic data can reach strong levels of evidence in germline variant interpretation in DD and enable reclassification of variants of uncertain significance. Together, these results establish somatic mutation data as a scalable and clinically actionable evidence source for germline variant interpretation in select DD genes.

**Graphical abstract:** (Generated using FigureLabs)

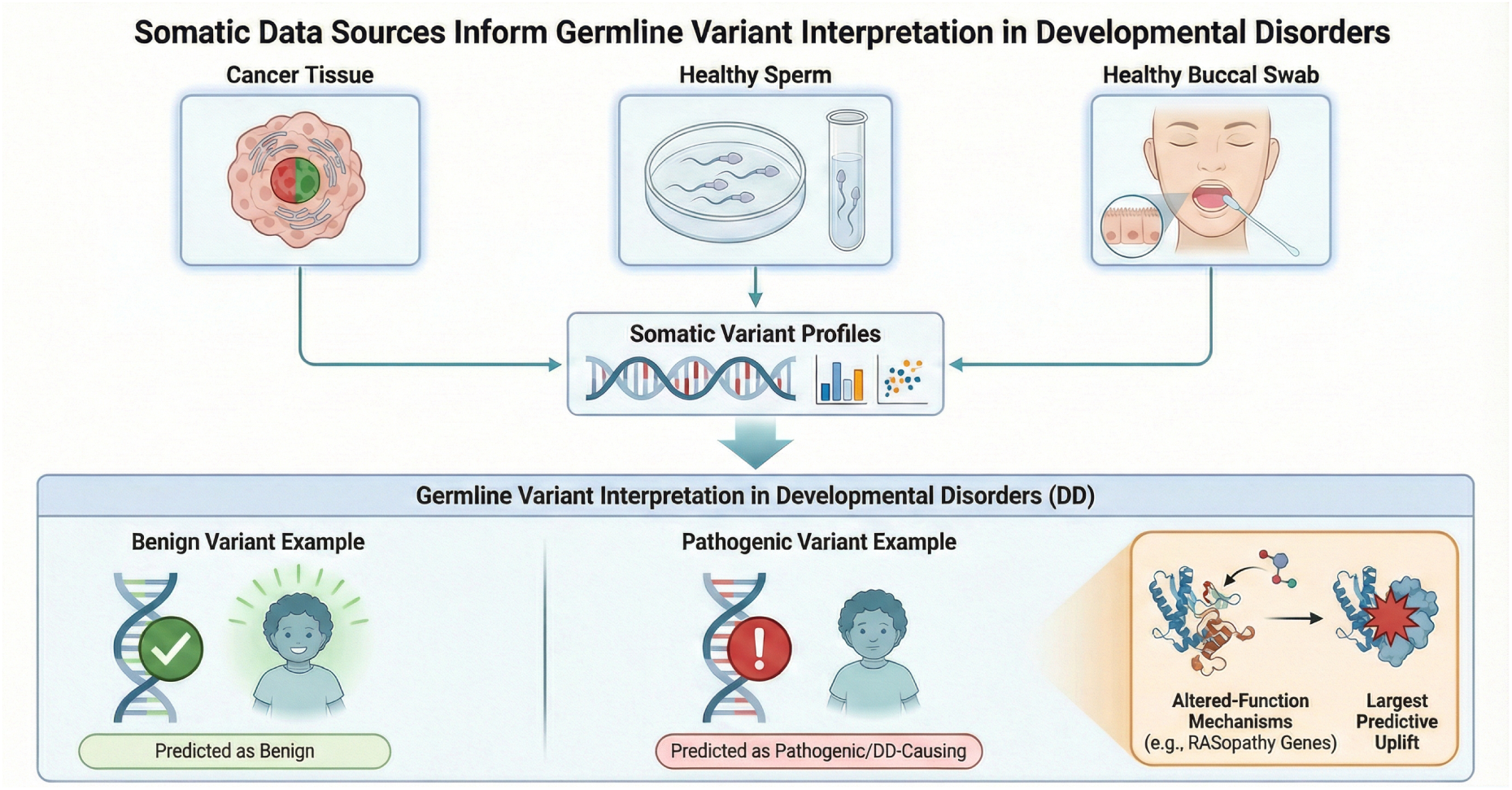

## 1 Introduction

Accurate prediction of variant impact remains a bottleneck for clinical interpretation of rare germline variation in developmental disorders (DD). Missense variants are particularly problematic: unlike truncating variants, their molecular consequences are highly context-dependent and cannot easily be inferred from sequence alone. The scale of this challenge is evident in ClinVar, where nearly half of curated variants are currently classified as variants of uncertain significance (VUS), with missense variants comprising the bulk of unresolved cases [1].

Compounding this difficulty is the difference between loss-of-function (LOF) and altered-function (AltFunc) disease mechanisms. AltFunc variants, encompassing gain-of-function and dominant-negative effects, are widespread among dominant DD genes, particularly in signalling pathways such as RAS–MAPK and PI3K–MTOR. The tools most widely used for variant interpretation are poorly suited to detecting AltFunc variants. Computational effect predictors (CEPs) model amino acid constraint or protein stability, features that predict LOF well but are poor at predicting AltFunc pathogenicity [2, 3]. Most scalable functional studies (Multiplexed Assays of Variant Effect, MAVEs) rely on cell viability or abundance readouts that capture LOF effects; assays with gain-of-function readouts exist for only a small fraction of DD genes. There is therefore a pressing need for additional evidence sources that can specifically discriminate pathogenic from benign variation in AltFunc genes.

One underused source of evidence is somatic variation: for some DD genes, the same aminoacid changes that are pathogenic in the germline are also observed as recurrent mutations under positive selection in somatic tissues as cancer driver variants. The mechanistic basis for germline–somatic overlap is well established in the RASopathies. The vast majority of activating substitutions observed in cancer are also observed as constitutional germline variants, and biochemical studies confirm that germline RASopathy mutations and their somatic counterparts act on the same functional residues through the same mechanistic perturbations [4]. Only the most strongly activating substitutions (e.g. *BRAF* p.Val600Glu) are absent as constitutional variants in viable individuals, consistent with the hypothesis that highly proliferative mutations are lethal when present constitutionally, while milder activating changes at the same functional residues produce viable developmental phenotypes [5]. Germline and somatic selection therefore act on largely overlapping substitutions at shared functional codons, making codon-level aggregation of somatic counts a natural strategy for leveraging cancer data in germline variant classification.

Large-scale analyses of cancer mutation hotspots show that recurrent somatic variants are enriched for pathogenic germline variants across diverse germline phenotypes, supporting the use of somatic recurrence as evidence in germline variant classification [6]. The ACMG/AMP guidelines provide the standard framework for germline clinical variant interpretation, and have been formalised as a quantitative system by modelling their evidence codes as likelihood ratios within a Bayesian classification framework, in which each evidence criterion is assigned a calibrated odds of pathogenicity and criteria are combined multiplicatively to yield a posterior probability of pathogenicity [7]. This quantitative approach enables evidence of different types and strengths to be integrated on a common scale, and provides a principled basis for deriving gene-or variant-class-specific thresholds. Somatic variation data have already been incorporated into variant interpretation frameworks for germline tumour predisposition genes [8]. For instance, the ClinGen Germline–Somatic Working Group has proposed guidance for applying PM1 (ACMG variant classification code for’located in a mutational hotspot’) in tumour predisposition genes using counts from CancerHotspots.org: PM1 is suggested when the exact amino-acid change has at least 10 tumour samples reported, and PM1-supporting when the count is 2–9 [9]. Recognising that the generic ACMG/AMP rules require contextual adaptation, ClinGen has established Variant Curation Expert Panels (VCEPs) for specific gene groups and associated disorders. Each VCEP is tasked with defining which evidence criteria apply to their gene set, at what strength, and with what quantitative thresholds. For example, the RASopathy VCEP has curated gene-specific rules for the RAS-MAPK pathway genes underlying Noonan syndrome and related conditions [10]. There is increasing evidence that many normal healthy somatic tissues are mosaics of expanding clones that have acquired mutations conferring a selective advantage. Such widespread somatic selection has been demonstrated in sun-exposed skin [11], oesophageal epithelium [12], buccal epithelium [13], and sperm [14]. These studies show that recurrent, positively selected mutations are a general feature of normal tissue biology, not restricted to cancer. Recent technical advances, including highly accurate duplex sequencing approaches such as NanoSeq [15], now enable ultra-deep detection of rare somatic variants. Unlike cancer samples, which often contain a small number of dominant driver mutations, healthy tissue samples typically harbour many independent expanding clones, so sequencing a single sample can reveal a broad spectrum of positively selected mutations and provide a rich resource for germline variant interpretation.

Despite these advances, somatic data have not been systematically benchmarked for germline variant interpretation in DD, and somatic sequencing data from healthy tissues have not been evaluated for germline pathogenicity prediction potential. Here, we test whether incorporating somatic evidence improves variant impact prediction in DD. We find 145 genes where somatic data can predict germline pathogenicity. The greatest clinical utility is in AltFunc mechanism genes, particularly the RASopathies, where somatic evidence achieves likelihood ratios consistent with strong-level evidence under established variant interpretation frameworks.

## 2 Methods

### 2.1 Gene mechanism classification

Developmental disorder (DD) genes were obtained from DDG2P ([16, 17]; accessed April 16, 2025). Analyses were restricted to genes with monoallelic DDG2P entries (autosomal or X-linked heterozygous) unless otherwise stated. Cancer gene mechanism annotations were obtained from the COSMIC Cancer Gene Census (CGC) (v98; [18]).

AltFunc genes were defined as the intersection of DDG2P monoallelic genes annotated with a probable altered-function mechanism (’altered gene product structure’ variant consequence) with COSMIC CGC oncogenes for which missense was a documented mutation type. The missense filter on COSMIC oncogenes was applied to exclude genes driven primarily by gene fusions or copy-number amplification. LOF genes were defined as the intersection of DDG2P monoallelic genes with a probable loss-of-function mechanism (’absent gene product’ variant consequence) with COSMIC CGC tumour suppressor genes.’Other’ DD-somatic selection genes comprised monoallelic DDG2P genes not meeting AltFunc or LOF criteria that nonetheless overlapped with either (i) somatic-selection genes identified in published sperm or buccal NanoSeq studies [14, 13] or (ii) the broader COSMIC CGC.

RASopathy genes were defined as those determined by the ClinGen Variant Curation Expert Panel (VCEP) [10] (*BRAF*, *HRAS*, *KRAS*, *MAP2K1*, *MAP2K2*, *NRAS*, *PTPN11*, *RAF1*, *RIT1*, *SHOC2*, *SOS1*, and *SOS2*). An AltFunc-minus-RASopathy subset was defined to enable analyses that isolate RASopathy-specific signal from the broader altered-function group.

Additional gene groups were defined from panels and pathways as follows:

- RASopathy genes defined by UK PanelApp (v1.86).
- Segmental overgrowth genes defined by the union of two UK PanelApp panels: Segmental overgrowth disorders – Deep sequencing (panel 98 v3.16) and Neurological segmental overgrowth (panel 736 v3.1).
- Macrocephaly/Megalencephaly panel from Australia PanelApp (accessed 23/3/26).
- Brain malformations VCEP group gene list: *AKT3*, *PIK3CA*, *PIK3R2*, *MTOR*, *DEPDC5*.
- Dominant macrocephaly genes were defined as genes from the Macrocephaly/Megalencephaly panel above that intersect with monoallelic DDG2P genes.
- PI3K–MTOR pathway genes were defined from WikiPathways (WP3844; [19]; accessed 23/3/26).
- RAS–MAPK pathway genes were defined from the KEGG hsa04010 MAPK signaling pathway list ([20]; accessed 23/3/26).

### 2.2 Germline variant datasets

Pathogenic, uncertain, and benign germline variants in DDG2P genes were obtained from ClinVar (accessed 6 February 2025, release 2025-02-06 [1]). De novo mutations (DNMs) from a cohort of 31,058 parent-proband trios were used as an alternative pathogenic variant list [21]. These lists were restricted to missense variants in Matched Annotation from NCBI and EMBL-EBI (MANE) Select transcripts within DDG2P genes, as called by bcftools csq [22]. ClinVar submissions were filtered to those with at least one germline entry, and the number of submissions to ClinVar was re-counted using only independent germline submissions to minimise somatic contamination.

Putative benign variants were obtained from the UK Biobank study [23], filtered to missense variants in DDG2P gene MANE select transcripts, with quality control and annotation performed as outlined in Gardner et al.[24]. No allele frequency filters were applied as the majority of missense variants were rare. Any variant annotated as ClinVar (likely) pathogenic or matching a DNM was removed.

The ClinVar and DNM datasets were prepared in both *unique* form (one row per distinct variant) and *count-weighted* form (one row per ClinVar germline submission or one row per proband with a DNM).

All variants were annotated with consequence, transcript, and codon position using bcftools csq, with a panel of computational effect predictors (CEPs) from dbNSFP version 5.2a [25, 26, 27, 28, 29, 30], and with somatic counts (see below).

### 2.3 Somatic mutation datasets and feature extraction

Each germline variant was annotated with four classes of somatic evidence: (i) COSMIC (v98) [18] variant-level counts from combined whole-genome and targeted sequencing and codon-level missense counts; (ii) sperm NanoSeq variant counts and codon-level missense counts [14]; (iii) buccal NanoSeq variant counts and codon-level missense counts [13]; and (iv) cancer hotspot status (Chang et al. hotspots v2, excluding frameshift-and splice-dominated codons [9]). Variants absent from a given somatic resource were assigned a count of zero.

The sperm NanoSeq and COSMIC data are counts from a mixture of panels and exome sequencing, whilst for the buccal data only panel data was available. If a gene was not covered in the buccal NanoSeq panel the area under the receiver operating characteristic curve (AUC) value was computed as NA (missing) rather than 0.5 (uninformative with exclusively zero values). For the sperm nanoseq data some genes only had hotspot coverage rather than full gene coverage in early iterations of their panel design. These data were removed to prevent circularity (high sperm NanoSeq counts seen in known hotspots).

### 2.4 Benchmarking framework for pathogenicity prediction performance

Pathogenic and benign variants were combined into two primary comparisons: ClinVar pathogenic versus UK Biobank (ClinVar datasets), and DNMs versus UK Biobank (DNM datasets). Each comparison was prepared in unique and count-weighted form.

For each gene group or individual gene and for each of the dataset forms described above, the performance of somatic data features and CEPs was measured. This was done using receiver operating characteristic (ROC) analysis (with 95% bootstrap confidence intervals), area under the precision–recall curve (AUPRC), sensitivity, specificity, precision, recall, accuracy, balanced accuracy, F1, and Matthews correlation coefficient (MCC). Where a fixed threshold was required, these were computed at the threshold maximising MCC among midpoint candidates between adjacent unique score values (capped at 500 threshold candidates). These metrics were only calculated if there were at least 10 unique pathogenic and benign variants available in the given gene or gene group with scores for the relevant feature.

Logistic regression models were trained to classify variants as pathogenic or benign using one CEP combined with log_1+_-transformed somatic evidence features. The model formula was:

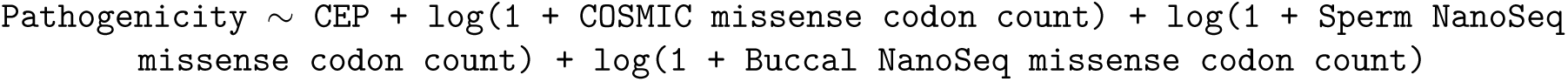

Three CEPs were evaluated in separate models: REVEL, AlphaMissense, and ESM1b. Training used 10-fold cross-validation repeated 10 times with random down sampling to address class imbalance. Models were evaluated on held-out test data (30%) by AUC. Statistical improvement over a CEP-only model was assessed with the DeLong test. Models were trained and evaluated across all gene group × dataset × CEP combinations. If count-weighted data was used, the train test split was performed on the unique form data before expanding to count-weighted data so as to expand each partition independently and prevent information leakage across splits.

When calculating models for individual genes, there were usually a small number of unique variants available, too few for a train-test split. Therefore, a leave-one-gene-out scheme was applied within each gene group. For each gene *g* in a given gene group, an logistic regression model was trained on all other genes in the group and evaluated on variants from gene *g*. Genes were included only if they contributed at least 10 unique benign and 10 unique pathogenic variants. A CEP was excluded for a given gene if *>* 20% of pathogenic variants had missing values. Model training used the same 10×10 repeated cross-validation with downsampling as above.

Positive likelihood ratios (LR+) and negative likelihood ratios (LR−) were calculated for 40 binary rules applied to five features: COSMIC count, COSMIC missense codon count, sperm NanoSeq count, sperm NanoSeq missense codon count, and cancer hotcodon status. For each rule, sensitivity and specificity were computed from 2×2 contingency tables against pathogenic versus benign variants, with 0.5 added to all cells when any cell was zero. Standard errors on the log scale were propagated to derive 95% confidence intervals by exponentiation. Analyses were stratified by gene group and repeated across four dataset forms (ClinVar unique, ClinVar count-weighted, DNM unique, DNM count-weighted). LR+ lower confidence intervals were mapped to ACMG evidence strengths using the Tavtigian Bayesian framework [7]: LR+ ≥ 2.08 (Supporting), ≥ 4.33 (Moderate), ≥ 18.7 (Strong), ≥ 350 (Very Strong).

### 2.5 Comparison with Multiplexed Assays of Variant Effect

For the 195 monoallelic DDG2P genes that were also involved in somatic selection (the AltFunc, LOF, and Other group genes described above), the literature was searched for MAVE studies using Edison Scientific’s Kosmos platform [31] (Edison Scientific, San Francisco, CA) and a query of MAVE-db [32]. Candidate studies were filtered to those that had full protein coverage for single nucleotide variants (rather than single domains), and to those that measured a relevant mechanism of pathogenicity. For example, an abundance assay would not be suitable to benchmark performance at distinguishing gain of function pathogenic germline variants from benign variants. A record of MAVE studies screened and rejected is available in Supplementary Table 3. Suitable MAVE studies were identified in *PTPN11*, *PTEN*, *DDX3X*, and *HRAS* [33, 34, 35, 36, 37, 38]. For *DDX3X*, the probabilistic NDD clinical classification score (probability of non-function) from a deep mutational scan was used, taking the maximum score across assay replicates per amino acid change.

If a suitable MAVE assay was available, the MAVE data was benchmarked in the same way as somatic data (see above), with the same benign and pathogenic variant lists. MAVE performance (AUC) was compared to the germline predictive performance of CEPs and somatic features.

### 2.6 Variant reclassification using somatic evidence

ClinVar missense variants in RASopathy genes were classified under the ACMG/AMP framework using a Bayesian point-based approach (prior pathogenic probability = 0.10) using codes that were feasible to automate. Where possible, application rules from the RASopathy VCEP were followed [10].

Allele-frequency cutoffs and residues used for PM1 followed the RASopathy VCEP guidelines. GnomAD version 4.1 [39] population maximum allele frequencies (popmax AF) were used. Likelihood ratios were PS = 18.7, PM = 4.33, PP = 2.08, VSb = 1/350, BS = 1/18.7, and BP = 1/2.08.

**Table 1:** ACMG/AMP evidence rules and likelihood ratios used for automated Bayesian classification of missense variants in RASopathy genes.

| Code | Strength | Application | LR applied |
| --- | --- | --- | --- |
| **Pathogenic evidence** |  |  |  |
| PS1 | Strong | Same amino acid change as established pathogenic variant (ClinVar $\geq 2\star$, exact nucleotide match) | 18.7 |
| PS2 | Moderate | De novo (for ClinVar variants, the de novo status described in ClinVar was used at moderate strength because of lack of maternity and paternity confirmation information) | 4.33 |
| PM1 | Moderate | Located in mutational hotspot or critical functional domain (codons defined by the RASopathy VCEP) | 4.33 |
| PM2 | Supporting | Absent from gnomAD | 2.08 |
| PP2 | Supporting | Missense variant in gene with low benign missense variation rate: applied to *BRAF*, *MAP2K1*, and *PTPN11* (GnomAD missense z score $> 3.09$) | 2.08 |
| PP3 | Supporting | Computational evidence: REVEL $\geq 0.70$ | 2.08 |
| **Benign evidence** |  |  |  |
| BA1 | Stand-alone | gnomAD popmax allele frequency $> 0.0005$ | 1/350 |
| BS1 | Strong | gnomAD popmax allele frequency 0.00025–0.0005 | 1/18.7 |
| BP4 | Supporting | Computational evidence: REVEL $\leq 0.30$ | 1/2.08 |
| **Bayesian framework parameters** |  |  |  |
| Prior probability |  | 0.10 (prior odds = 0.111) |  |
| Classification thresholds | | Pathogenic $\geq 0.99$; Likely pathogenic $\geq 0.90$; Likely benign $< 0.10$; Benign $\leq 0.001$ | |

### 2.7 Software

All analyses were conducted in R (≥4.3). Key packages included tidyverse, pROC, caret, and PRROC. Reports were generated using R Markdown knitted to HTML via Pandoc (v3.4).

## 3 Results

### 3.1 Large overlap between DD genes and genes driving somatic selection

The large overlap between genes that cause DD and those associated with selection in cancer is well documented [40]. Across 970 dominant DD genes [17], 195 drive somatic selection in at least one context: cancer (as curated in the COSMIC Cancer Gene Census [18]), buccal epithelium [13], or sperm [14] (Supplementary Figure S1).

A subset of these genes plausibly act through the same underlying mechanism in both germline DD and somatic selection (both altered function or both loss of function). We hypothesised that these “mechanism-matched” genes would have the greatest utility of somatic data in germline variant impact prediction. We defined gene groups by integrating developmental-disorder mechanism annotations from DDG2P [17] with cancer mechanism annotations from the COSMIC Cancer Gene Census [18] (table 2). Of the 195 dominant DD genes implicated in somatic selection, 40 were identified as altered function in both DD and cancer, 63 as both loss of function (with an overlap of 3 genes meeting the criteria for both groups), and the remainder (’Other’) had missing annotations or mismatched mechanisms.

**Table 2:** Gene groups defined by mechanism annotations in developmental disorders (DD) and cancer.

| Gene group | Mechanism (DD) | Mechanism (cancer) | Number of genes |
| --- | --- | --- | --- |
| AltFunc | Altered function | Oncogene | 40 |
| LOF | Loss of function | Tumour suppressor gene | 63 |
| Other DD-somatic genes |  |  | 95 |

### 3.2 DD-causing variants overlap with somatic selection variants

Two sets of pathogenic variants were used to quantify overlap between DD-causing mutations and somatic selection. First, we extracted ClinVar pathogenic/likely pathogenic germline missense variants in DD genes (90,696 submissions; 35,886 unique variants across 2,087 genes) [1]. Second, we analysed published missense de novo mutations (DNMs) from 31,058 parent–offspring trios with developmental disorders [21], comprising 8,674 variants across 1,821 DD genes. As a benign control, we used missense variants from UK Biobank participants in the matched gene set [41] (1,193,326 putative benign variants across 2,403 DD genes).

Because relative mutation frequency informs assessment of concordance between DD variation and somatic selection, we analysed pathogenic variants in both “count-weighted” and “unique” form: count-weighted data includes each variant once per observation, whereas unique-form data includes each distinct variant once. Unless otherwise stated, results show count-weighted ClinVar pathogenic variants; key findings were replicated using unique-form ClinVar variants and DNMs (Supplementary tables S1, S2, S4, S5, S6).

Mechanism-matched altered-function genes show the clearest mirroring between germline and somatic mutation patterns (e.g. *MAP2K1*, *RIT1*, and *KRAS*; Figure 1), though relative frequencies of clusters of proximal mutations can differ between contexts suggesting tissue-specific selection pressures. In genes with discordant DD and somatic mechanisms, ClinVar may include non-DD phenotypes—for example, *SMAD4* altered-function variants associated with Myhre syndrome overlap with sperm sequencing variants[42], while cancer patterns more closely resemble germline juvenile polyposis variants present in ClinVar but absent from DD cohort DNMs (Figure 1).

**Figure 1:**
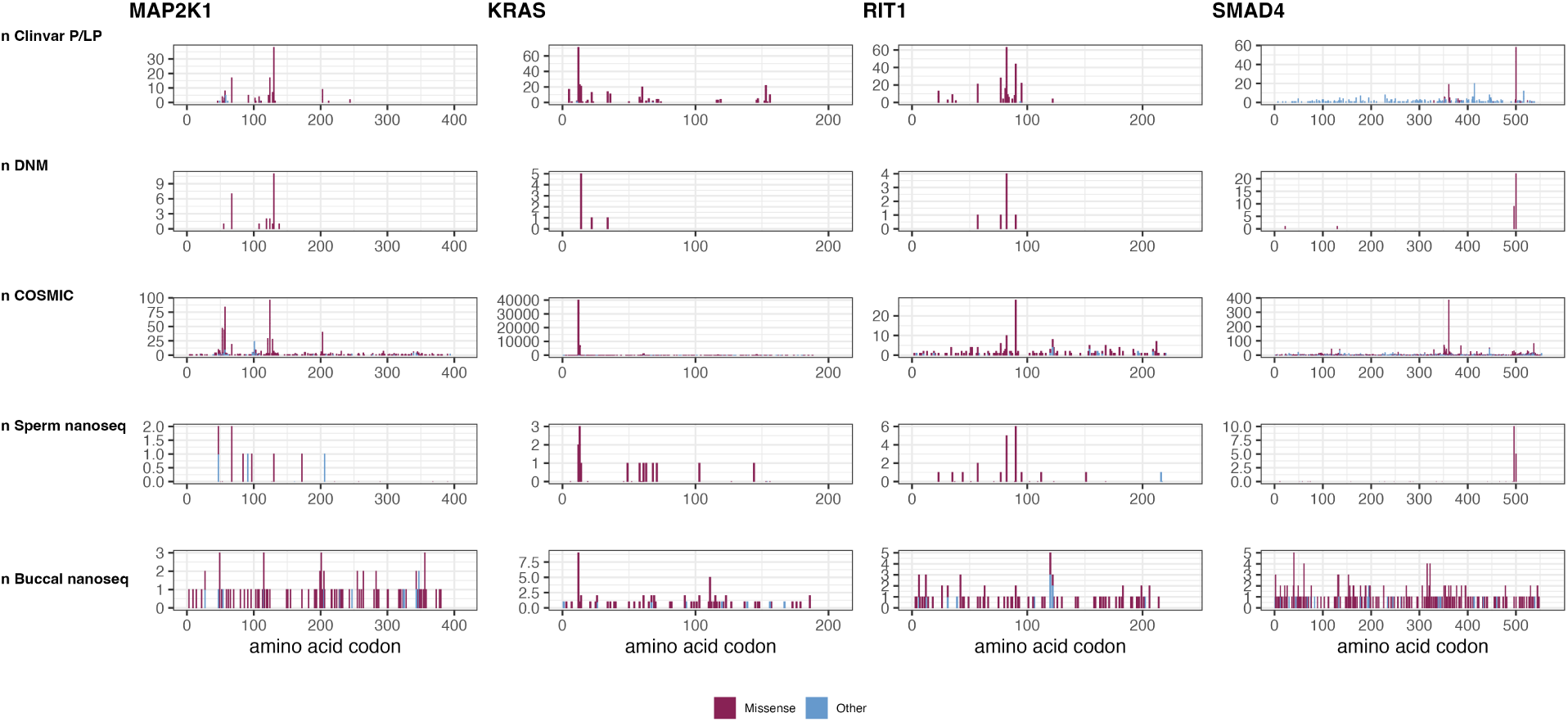
Mutation frequency and distribution across example proteins, illustrating mirrored (MAP2K1, KRAS, RIT1) and discordant (SMAD4) patterns between DD and somatic selection datasets

### 3.3 Somatic data predicts germline pathogenicity in DD genes

We assessed how well somatic data features perform as germline pathogenicity predictors in each of the gene groups defined above. From each somatic data source (cancer, sperm and buccal tissues) we evaluated the performance of the raw count (number of times the germline variant is seen in the somatic database), and we also pooled missense variants at a codon level to create a’missense codon count’. This improves the sensitivity and performance of these sparse somatic data (see Figure 2a) (Supplementary table S1). This is because DD-causing variants can be at the same codon, but cause a different amino-acid change, from somatic hotspot mutations.

**Figure 2:**
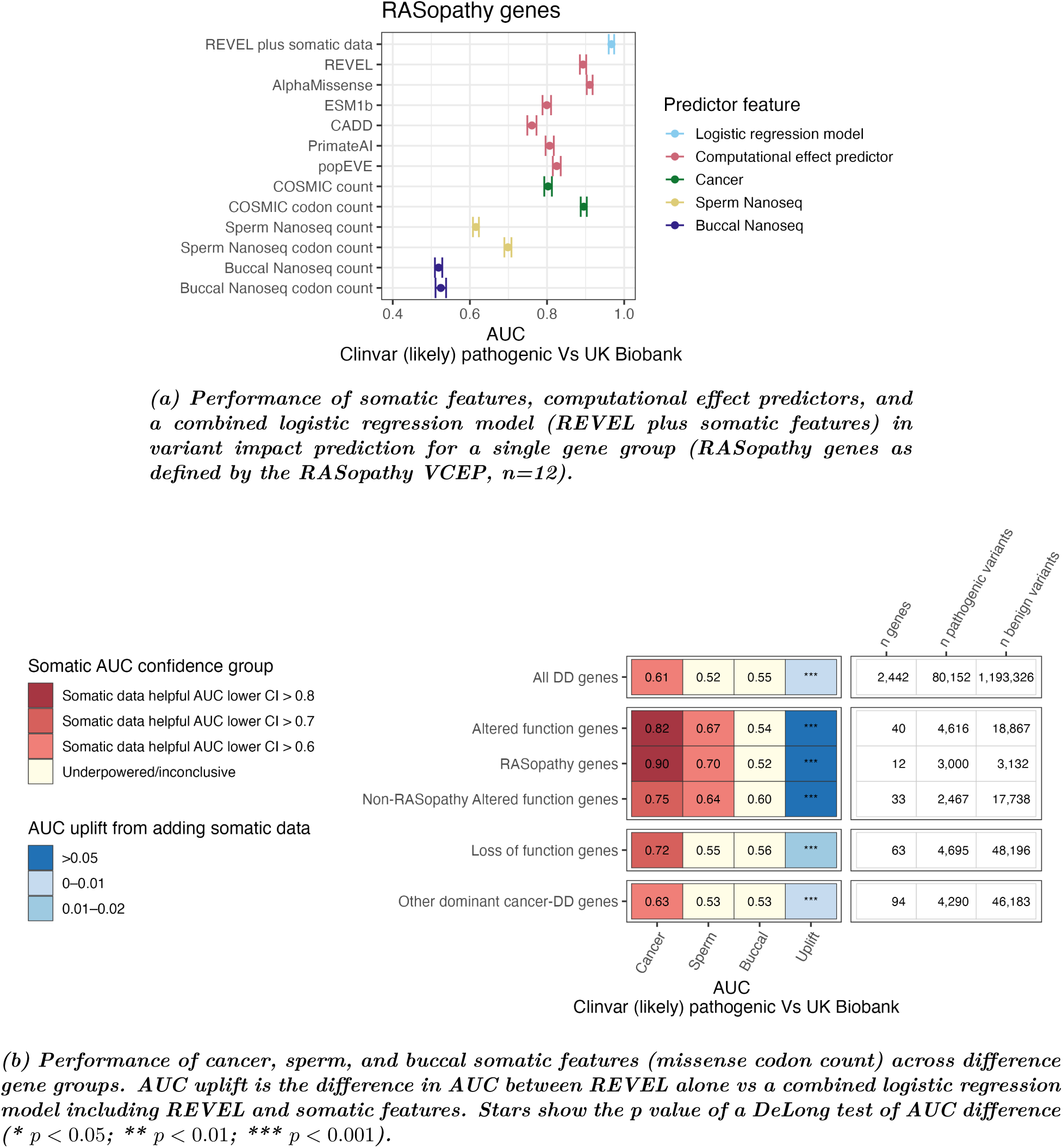
Predictive performance of somatic-selection features for germline pathogenicity classification.

In RASopathy genes the COSMIC missense codon count feature achieves an AUC of 0.90 (95% confidence interval (CI) 0.89–0.90) to distinguish benign from pathogenic germline variants, comparable to widely used CEPs such as REVEL (AUC 0.89, 95% CI 0.89–0.90) [26]. In contrast, the sperm NanoSeq codon missense counts shows lower, but still informative, discrimination (AUC 0.70, 95% CI 0.69–0.71), despite the data being sparse (currently from only 81 sperm samples). The buccal sequencing data tends to have poor discriminatory power in all but a few DD genes (overall AUC for RASopathy genes 0.52, 95% CI 0.51-0.54). If the RASopathy genes are removed from the altered function gene group, the remaining genes (’Non-RASopathy altered function genes’) still show good predictive performance of somatic mutation datasets (see Figure 2b), suggesting there are more altered function genes beyond the RASopathies in which these data could be used. Loss of function genes and non-mechanism matched (’other’) DD somatic genes showed more modest utility of somatic data (Figure 2b).

To look at individual genes rather than groups of genes, we restricted benchmarking to genes with at least 10 unique benign and 10 unique pathogenic variants (705 DD genes total for the ClinVar count-weighted data). This identified 145 genes for which somatic data can distinguish germline pathogenic from benign variants, defined by a somatic AUC lower confidence interval of *>* 0.6: 138 genes passed this threshold using the COSMIC missense codon count, 16 genes using the sperm NanoSeq missense codon count, and 16 using the buccal NanoSeq missense codon count (Supplementary table S2). The sperm NanoSeq data are likely to become more informative as additional samples are sequenced: AUC point estimates are frequently high, but confidence intervals are currently too wide to be classified as helpful under this metric.

### 3.4 Somatic data can out-perform or complement MAVEs and computational effect predictors

We next evaluated if somatic data adds predictive power in addition to existing germline variant impact prediction tools: computational effect predictors (CEPs), and functional data (MAVEs).

Among the 195 dominant DD somatic selection genes, 124 had sufficient variants for individual bench-marking (defined as ≥ 10 unique pathogenic and benign with REVEL score); somatic codon count alone (COSMIC, sperm, or buccal) exceeded the AUC performance of REVEL with non-overlapping confidence intervals in 8 genes (Supplementary table S2). Whilst it is rare that a somatic dataset alone is able to outperform a CEP, there are many more genes where combining somatic data with a CEP in a logistic regression model produces significantly better predictive power than the CEP alone (n=50 genes with DeLong p<0.05, combined AUC > REVEL AUC, Supplementary table S5). At a gene group level, this ‘diagnostic uplift’ is more obvious: the combined model significantly outperformed REVEL alone in the altered function gene group (AUC 0.95, 95 % CI 0.95–0.96 vs. REVEL alone AUC 0.90, 95 % CI 0.89–0.90; DeLong *p* = 3.0 × 10*^−^*^65^) and the RASopathy gene group (AUC 0.97, 95 % CI 0.96–0.97 vs. REVEL alone AUC 0.89, 95 % CI 0.89–0.90; DeLong *p* = 3.4 × 10*^−^*^20^) (Figure 2, Supplementary table S4).

The genes showing the greatest diagnostic uplift from a combined model compared to REVEL alone were those in which somatic data performs well but CEPs perform poorly. In genes where REVEL already achieves high discriminative performance, the potential for improvement is limited. For example in *BRAF*, REVEL achieves an AUC of 0.93 (95 % CI 0.90–0.95) and the combined model provides only a modest improvement (combined AUC 0.946, 95 % CI 0.93–0.97; ΔAUC = +0.021, DeLong *p* = 0.04). In contrast, for *PIK3CA* CEPs are weaker (REVEL AUC 0.74, 95 % CI 0.70–0.79). Here, somatic codon counts substantially outperform CEPs (COSMIC codon count AUC 0.97, 95 % CI 0.95–0.99) and the combined model yields the largest individual-gene uplift in our dataset (ΔAUC = +0.212, DeLong *p* = 3 × 10*^−^*^28^).

There were 22 genes where the combined model with somatic features significantly reduced performance relative to REVEL alone (DeLong p < 0.05, combined AUC < REVEL AUC); these were predominantly LOF and non-mechanism matched (’other’) genes. This highlights the importance of not using somatic data indiscriminately.

Massively parallel functional assays (MAVEs) provide an attractive source of quantitative evidence for germline variant interpretation, but MAVE data are not yet available for most DD genes and, where available, the functional readout may not match the relevant disease mechanism. Among the 195 dominant DD genes implicated in somatic selection, 4 have scalable MAVE data with good coverage of pathogenic DD variants and a functional readout that captures the same mechanism as pathogenic DD variation (literature review summarised in supplementary table S3, [33, 34, 35, 36, 37, 38]). MAVEs covering restricted protein regions containing too few DD pathogenic variants, or with an output that does not reflect the DD mechanism (e.g. a loss of abundance assay in an altered function DD gene) were not evaluated.

We compared somatic features and MAVE scores as discriminators of germline pathogenicity. For the altered function gene *PTPN11*, the COSMIC missense codon count outperformed MAVE data. For other genes, the MAVE data had similar performance (*PTEN*, *HRAS*) or superior performance (*DDX3X*) to the COSMIC missense codon count (Figure 3a).

**Figure 3:**
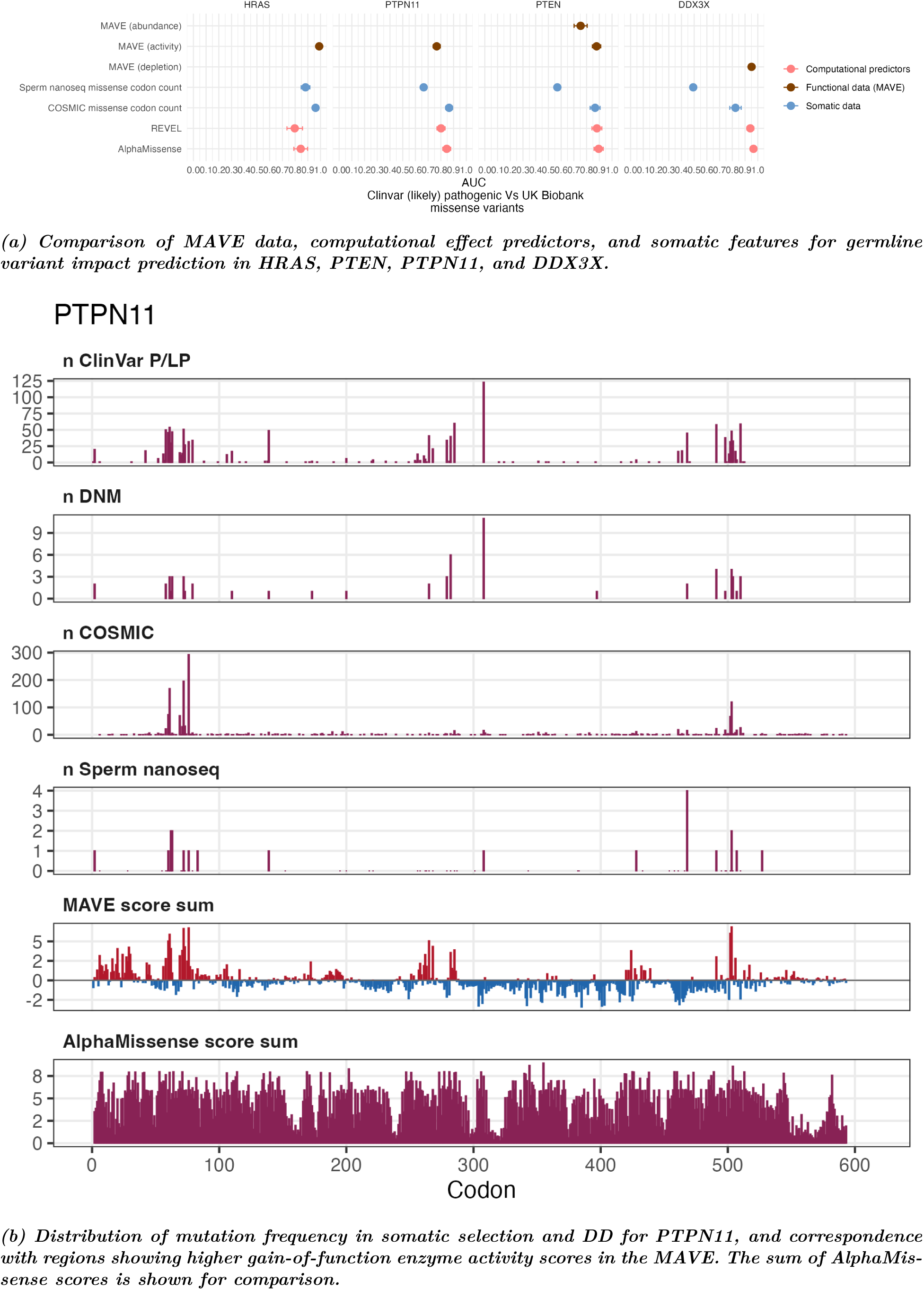
Somatic data can out-perform or riv1a1l MAVEs and computational effect predictors.

For *PTPN11*, somatic hotspots and MAVE enrichment peaks line up with the germline pathogenic variants (Figure 3b) whilst AlphaMissense scores fail to highlight the gain of function hotspot regions, presumably because they also have high scores for loss of function variants.

### 3.5 Clinically meaningful grouping of genes for application of somatic evidence

Within the gene groups defined above according to mechanism, we observed heterogeneity in the utility of different somatic data sources (Supplementary Figure S2). In contrast, the RASopathy gene group (defined by the ClinGen RASopathy variant curation expert panel, VCEP) show uniformly high utility of somatic data (see Figure 4).

**Figure 4:**
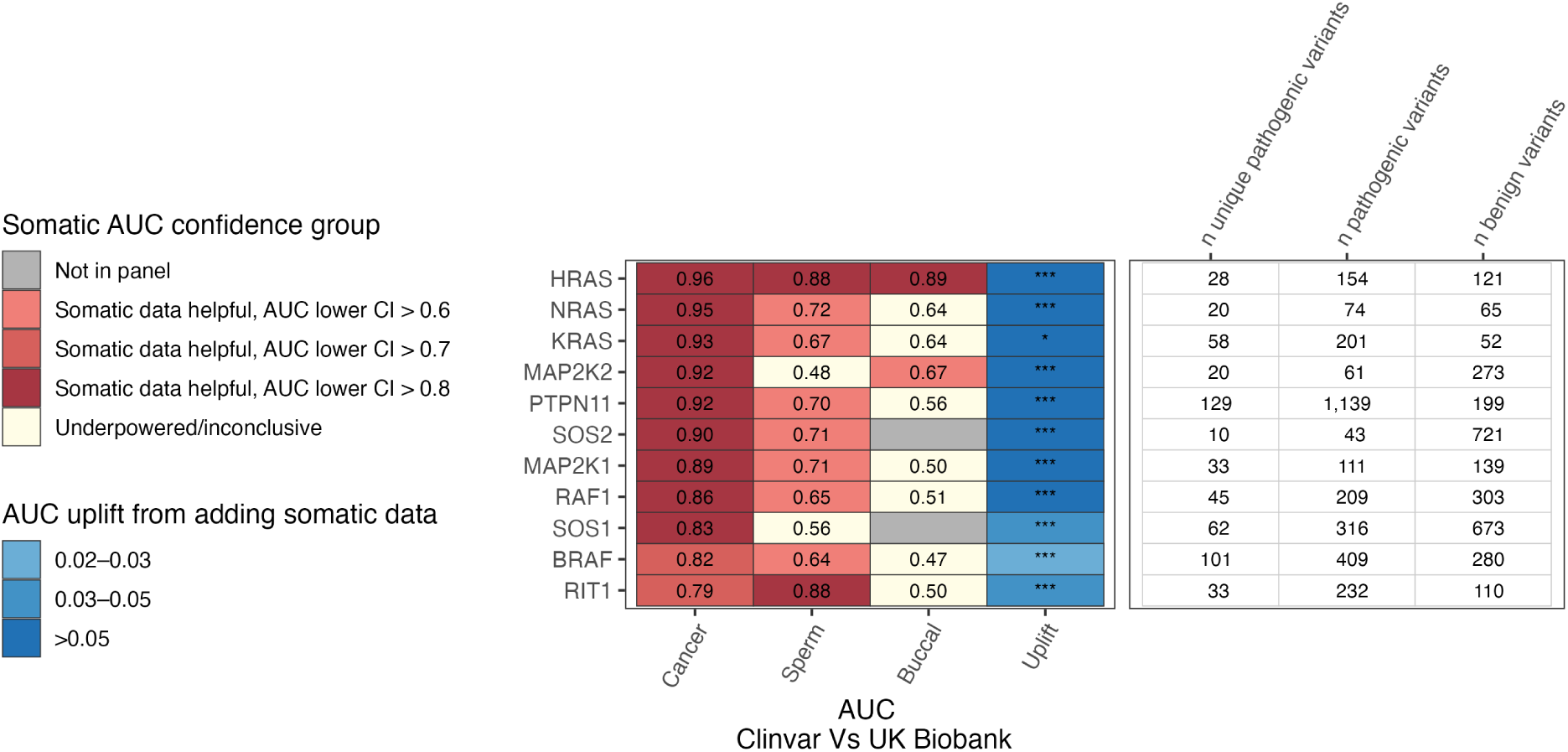
Utility of somatic data in the RASopathy gene group (ClinVar count-weighted-form benchmarking dataset). 11 of the 12 VCEP RASopathy genes are shown (SHOC2 excluded due to fewer than 10 unique pathogenic ClinVar variants)

One approach to clinical application of these data could be to apply somatic data in variant interpretation in any gene that passes a benchmarking threshold (e.g. lower CI of AUC *>* 0.6 for a somatic feature). However, some of these genes lack power to assess individually. Additionally, modifications to variant interpretation guidelines are usually published for mechanistically and clinically homogeneous groups of genes (largely via Variant Curation Expert Panels - VCEPs). With this in mind, we have explored ways to group DD genes that have the highest utility of somatic data. Many of the top scoring genes belong to the RAS–MAPK or PI3K–MTOR pathways, or are implicated in overgrowth-like phenotypes such as segmental overgrowth or macrocephaly (Table 3).

**Table 3:** Utility of somatic data across gene groups defined by clinical criteria or molecular pathways, summarised as the number (%) of genes whose COSMIC missense codon count AUC lower confidence interval exceeds different thresholds.

| Category | *n* genes | *n* (%) with AUC lower CI > 0.6 | *n* (%) with AUC lower CI > 0.8 |
| --- | --- | --- | --- |
| VCEP: Brain malformations (*PIK3CA/MTOR*) | 4 | 4 (100%) | 3 (75%) |
| VCEP: RASopathies | 12 | 12 (100%) | 9 (76%) |
| Panel: RAS-MAPK (UK PanelApp) | 18 | 16 (89%) | 12 (67%) |
| Panel: Segmental overgrowth (UK PanelApp) | 7 | 6 (86%) | 4 (57%) |
| Phenotype: Dominant macrocephaly | 57 | 31 (54%) | 16 (28%) |
| Pathway: PI3K-MTOR (WikiPathways; [19]) | 8 | 8 (100%) | 7 (88%) |
| Pathway: RAS-MAPK (KEGG) | 18 | 10 (56%) | 8 (44%) |

Defining gene groups according to clinical features (variant interpretation working group gene lists, diagnostic panels, or phenotypes), rather than molecular pathways (e.g. WikiPathways or KEGG terms) produces a more meaningful list of genes in which somatic data can be benchmarked for clinical diagnostic application (see Table 3). For example, all of the genes defined by the RASopathy VCEP group, which cause the relatively homogenous dominant Noonan-syndrome phenotypes, have high somatic utility. In contrast, the genes in the RAS–MAPK KEGG pathway are more clinically heterogenous and can even be associated with growth restrictive phenotypes (e.g. *IGF1R*) or no known DD phenotype (e.g. *EGFR*, *PDGFRA*, *PDGFRB*).

### 3.6 Addition of somatic data resolves variants of uncertain significance in RASopathy genes

We quantified how somatic signals map onto ACMG/AMP-style evidence strength using likelihood ratios (LRs) (Supplementary table S6). Hotcodon status from CancerHotspots.org produced the highest LRs (e.g., for altered-function genes using ClinVar count-weighted data, LR 91.2 [71.3–117]), but with limited sensitivity. Fewer than 1% of RASopathy variants annotated in ClinVar as VUS or conflicting interpretations fall within cancer mutation hotcodons, limiting the utility of this feature for resolving uncertain classifications. For higher sensitivity, COSMIC missense codon counts provided a useful trade-off while still reaching supporting-to-strong evidence levels using the LR lower confidence bound; for example, in using ClinVar count-weighted data in RASopathy genes, a COSMIC missense codon count threshold of *>* 2 gives LR = 6.88 [6.22–7.60] (moderate evidence) at 75% sensitivity, while *>* 10 gives LR = 29.8 [22.7–39.0] (strong evidence) at 49% sensitivity (Figure 5a). These LRs are robust to the use of different pathogenic variant sources and using different forms of somatic data (i.e. count-weighted or unique) (Supplementary Figure S3).

**Figure 5:**
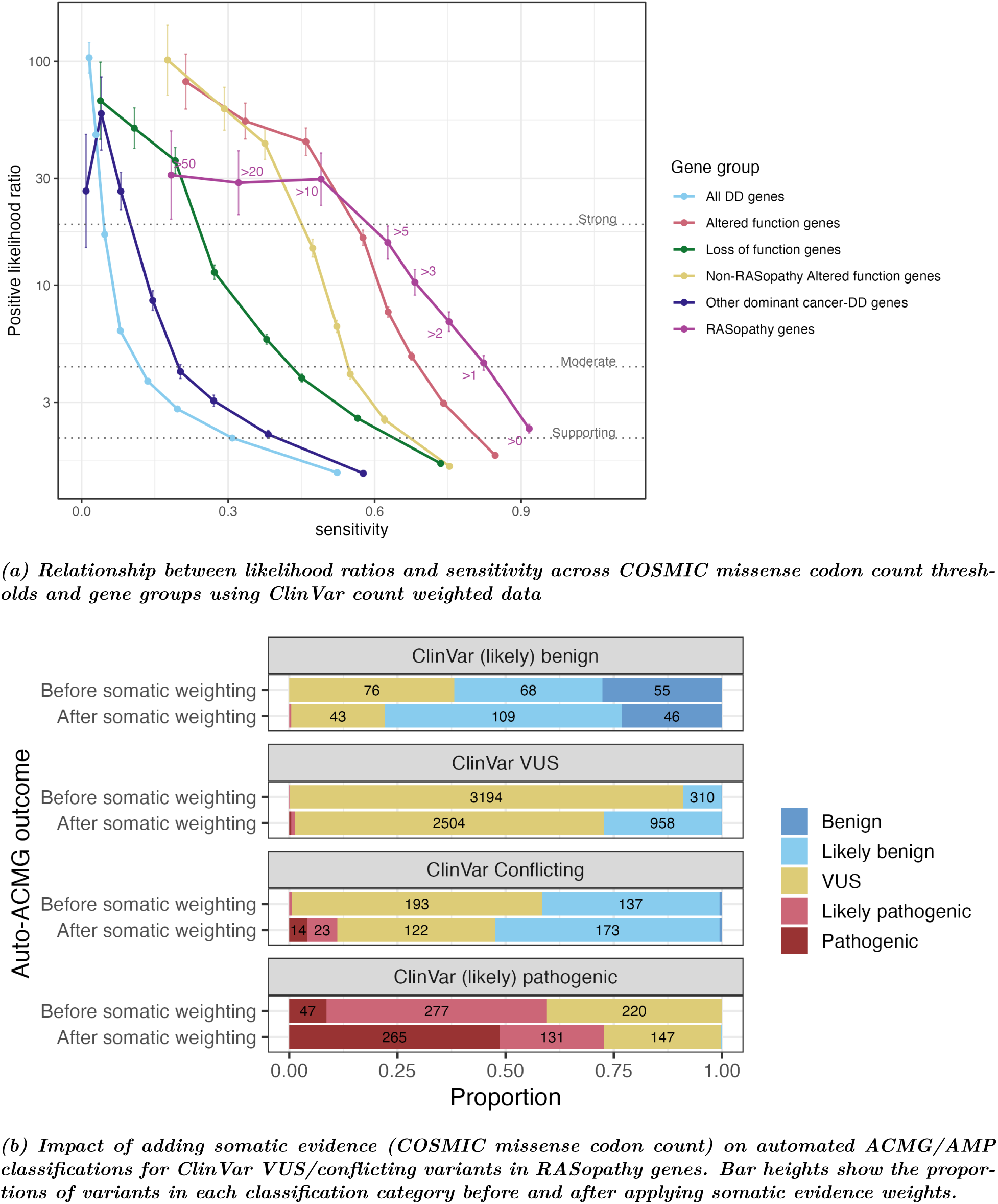
Calibration and application of somatic evidence for clinical use

To formalise the use of somatic evidence within an ACMG/AMP framework, we derived rules for applying COSMIC missense codon counts using the likelihood ratios estimated above, mapped to evidence strength thresholds under a Bayesian ACMG/AMP formulation [7]. To err on the side of caution, we used the lower confidence interval of each likelihood ratio when assigning evidence strength. We further compared likelihood ratios derived from four pathogenic variant lists before defining final cutoffs (see Figure S3).

This yielded the following somatic evidence rules for RASopathy genes:

- **Benign (supporting):** COSMIC missense codon count = 0.
- **Pathogenic (moderate):** COSMIC missense codon count *>* 2.
- **Pathogenic (strong):** COSMIC missense codon count *>* 10.

We annotated all ClinVar variants in RASopathy genes with additional evidence types commonly used in clinical interpretation, including computational effect predictor scores and gnomAD allele frequencies (see Methods). We then calculated Auto-ACMG scores for each variant using this publicly available evidence and the ClinGen RASopathy VCEP modifications to the ACMG/AMP framework. Variants were auto-scored both before and after adding somatic evidence at the weights listed above, treating somatic evidence as an independent evidence category.

Before the addition of somatic weighting, there were 193 variants marked as Conflicting and 3194 variants marked as VUS in ClinVar that our auto-ACMG code could not resolve. After the addition of somatic weighting, this reduced to 122 and 2504 respectively, a 22% reduction in the ClinVar VUS group and a 37% reduction in the ClinVar Conflicting group (Figure 5b). Roughly nine tenths of reclassifications were downward to (likely) benign, even with somatic weighting only applied at a conservative supporting level for benign evidence. In addition, a higher proportion of ClinVar (likely) pathogenic and (likely) benign variants could be correctly automatically classified if somatic data was included (59.6% to 72.8% and 61.8% to 77.9% respectively). This highlights the potential impact on reducing VUS burden in automatic variant interpretation workflows for RASopathy genes.

## 4 Discussion

This study demonstrates that somatic mutation data can be used to improve germline variant interpretation in an appreciable subset of developmental disorders (DD). By integrating somatic mutation recurrence metrics with germline variant datasets we show that somatic mutation data, especially codon-level aggregation of missense counts, provide predictive power that is (in some contexts) comparable to computational effect predictors and multiplexed functional assays. Importantly, somatic data adds independent information beyond these other evidence sources, improving classification performance when combined with them. We identify over 140 DD genes in which somatic data provides informative signal, with the greatest utility observed in altered-function genes where germline and somatic mechanisms are concordant. This is particularly evident in RASopathy genes, where somatic evidence can reach strong levels under ACMG/AMP-style frameworks.

Our work expands on prior descriptive studies demonstrating overlap between somatic mutation hotspots and germline pathogenic variants across diverse disease contexts [6, 40]. Existing guidelines from the ClinGen Germline-Somatic Working Group [8] have focused on using somatic data to inform germline variant interpretation in tumour predisposition genes, requiring recurrence of the exact amino-acid substitution in statistically significant ‘hot-codons’. While such criteria yield very high likelihood ratios, they are inherently sparse: only a small fraction of germline variants, particularly those classified as variants of uncertain significance (VUS), fall within these narrowly defined hotspots. Our results show that, in the context of DD, relaxing these criteria to incorporate codon-level aggregation substantially increases sensitivity while retaining clinically meaningful levels of evidence. This is biologically plausible, as germline and somatic selection often act on the same functional residues but favour different amino-acid substitutions depending on fitness constraints. Strongly activating mutations that dominate cancer (e.g. *BRAF* p.Val600Glu) may be embryonic lethal in the germline, whereas milder substitutions at the same codon produce viable but pathogenic developmental phenotypes (e.g. *BRAF* p.Val600Gly in cardiofaciocutaneous syndrome [43]). Consistent with this, we observe that missense codon counts outperform exact variant counts as predictors of germline pathogenicity.

Comparison with MAVE data highlights the subtleties of altered function mechanisms. In *PTPN11*, for example, the most common DD-associated mutation (p.Asn308Asp) has a modest MAVE score (0.1531) [34] and is not a major cancer hotspot in COSMIC, yet it is seen in sperm NanoSeq. Such discordance indicates that no single evidence type will capture the full spectrum of pathogenic gain-of-function variation. Somatic selection metrics from multiple tissue contexts, computational predictors, and MAVEs can therefore be most informative when interpreted as complementary, mechanistically distinct signals that can be combined. This is particularly important given that altered-function mutations remain challenging to predict using computational methods [2, 44] and are not well captured by the most scalable MAVE assays, which typically rely on loss-of-function or cell viability readouts.

Several biological factors likely explain the observed differences in performance between somatic data sources. Cancer datasets, such as COSMIC, provide the strongest and most broadly applicable signal. This is likely because of the scale (∼1.5 million samples), and because cancer represents a composite of many distinct selective environments across tissues. As a result, cancer data captures selection signals across hundreds of genes, increasing its utility for germline interpretation. In contrast, somatic data from normal tissues such as sperm and buccal epithelium reflect more restricted biological niches. Sperm sequencing, which captures selection in the male germline, shows clear relevance to DD genes, particularly those associated with paternal age-effect mutations. However, it is currently limited by sample size and data sparsity. Buccal epithelium data appears less informative for DD, likely reflecting weaker overlap between the selective pressures in this tissue and the mechanisms underlying developmental phenotypes. As larger datasets from normal tissues become available, particularly for germline-relevant contexts such as sperm, their utility is likely to increase. From a clinical and translational perspective, these findings have several implications. Achieving a molecular diagnosis in children with developmental disorders has well-established benefits for clinical management, prognosis, and family counselling [45]. Our results suggest that somatic data could be incorporated into ACMG/AMP-style guidelines for variant interpretation, but its optimal placement within the current guidelines remains to be determined. Existing guidance has positioned somatic recurrence within the PM1 (mutational hotspot) criterion for tumour predisposition genes [8]. However, somatic selection signals, especially those derived from codon-level aggregation, capture broader biological information than traditional hotspot annotations. In some respects, *in vivo* somatic data more closely resemble functional evidence derived from m*in vitro* proliferation-based assays. Somatic data may warrant consideration as a distinct evidence category, with careful attention to potential correlations with other evidence types. For example, proliferation-based functional assays and tumour recurrence are likely to be correlated and should not be treated as independent evidence without careful justification, whereas saturation genome editing assays measuring cellular viability, or biochemical MAVEs measuring enzyme activity, may provide mechanistically distinct information.

Although some somatic features achieve likelihood ratios consistent with strong evidence, their clinical adoption may follow a trajectory similar to that of computational predictors, which were initially applied conservatively at supporting strength despite strong statistical performance [46, 47, 48]. As additional benchmarking datasets and independent validations accumulate, there may be scope to increase the evidential weight assigned to somatic features.

This study has several limitations. First, the pathogenic variant datasets used for benchmarking have inherent constraints. ClinVar aggregates submissions across multiple phenotypes and may include misannotations or residual somatic variants despite filtering. In a minority of genes, the observed signal may be driven by non-DD phenotypes represented within ClinVar for nominally “DD” genes. For example, ClinVar entries for *LZTR1* are predominantly associated with schwannomatosis rather than Noonan syndrome. De novo mutation datasets only provide data for disorders with a strong de novo component to pathogenicity and not all de novo mutations are pathogenic. Power limitations arising from both variant datasets and somatic annotation resources likely lead to an underestimation of the number of genes in which somatic data is informative. Much of the somatic data comes from targeted panels rather than whole exome or genome sequencing. Our analyses are largely performed at the level of gene groups rather than individual genes. While this approach increases statistical power, it may obscure gene-specific differences in the relative utility of different somatic data sources. Although we note that the utility of computational effect predictors can also vary substantially between DD genes. As larger and more comprehensive somatic mutation datasets become available, particularly for normal tissue sequencing, additional genes are likely to reach thresholds of clinical utility.

Additionally, the integration of somatic data into existing variant interpretation frameworks remains an open challenge. The ACMG/AMP guidelines were not designed with this type of evidence in mind, and there is currently no consensus on how best to incorporate it. Somatic selection reflects biological fitness effects in specific cellular contexts. As a result, the relationship between somatic recurrence and germline pathogenicity is expected to vary by gene, mechanism, and tissue niche. The clinical community may therefore initially favour conservative application despite apparent strong statistical support. Realising the potential of these data will require not only the generation of large-scale somatic datasets, but also the development of infrastructure that makes them accessible and actionable at the point of clinical decision-making. Somatic variant databases will need to be continuously updated and structured in machine-readable formats (e.g. served via standardised APIs) that allow downstream integration into clinical tools such as DECIPHER [49].

Looking forward, the utility of somatic data for germline variant interpretation is likely to increase substantially with the continued expansion of both somatic sequencing datasets and curated germline variant resources. Improvements in sequencing technologies, including ultra-accurate methods such as duplex sequencing, will enable more comprehensive detection of low-frequency somatic variants across diverse tissues. Larger and better-annotated germline datasets, including longitudinal birth cohorts with detailed phenotypic follow-up, will provide a richer catalogue of both pathogenic and benign variants linked to well-characterised clinical outcomes, enabling more robust benchmarking and calibration of evidence strengths across the full spectrum of variant classification. Together, these advances will support the development of more precise, gene-specific models and facilitate integration into clinical practice, establishing somatic selection as a rich source of clinically actionable information for variant interpretation.

## 5 Data and code availability

All scripts used to generate the figures, tables, and claims in the text are available on GitHub: https://github.com/KatrinaAndrews/Somatic_germline

Access to the raw variant lists used for benchmarking is available by downloading ClinVar release 2025-02-06 [1], the de novo mutations from Kaplanis et al. [21], and the UK Biobank Resource.

**Supplementary table 1:** Benchmarking results for the use of somatic data in germline clinical variant interpretation for gene groups (including sensitivity, specificity, AUC, AU-PRC, etc., and including results using different pathogenic variant lists).

**Supplementary table 2:** Benchmarking results for the use of somatic data in germline clinical variant interpretation for individual genes.

**Supplementary table 3:** List of MAVE studies that were considered for inclusion in results section 4.4. **Supplementary table 4:** Performance of logistic regression models combining a computational effect predictor with somatic data and results for the AUC uplift this generates with DeLong p values (gene groups).

**Supplementary table 5:** Performance of logistic regression models combining a computational effect predictor with somatic data and results for the AUC uplift this generates with DeLong p values (individual genes).

**Supplementary table 6:** Likelihood ratios produced for different somatic feature cutoffs in different gene groups and with different pathogenic variant lists.

## 6 Acknowledgements

This research has been conducted using the UK Biobank Resource under application number 44165. This research was funded in whole, or in part, by the Wellcome Trust [Grant number 220540/Z/20/A]*. For the purpose of Open Access, the author has applied a CC BY public copyright licence to any Author Accepted Manuscript version arising from this submission. MT was supported by the NIHR Cambridge Biomedical Research Centre (NIHR203312). HVF is supported by the Wellcome Trust [226083/Z/22/Z] for PARADIGM (Primary Annotated Resources to Aid Diagnosis In Genomic Medicine).

## 7 Conflicts of interest

None.

## Supplementary Figures

**Figure S1:**
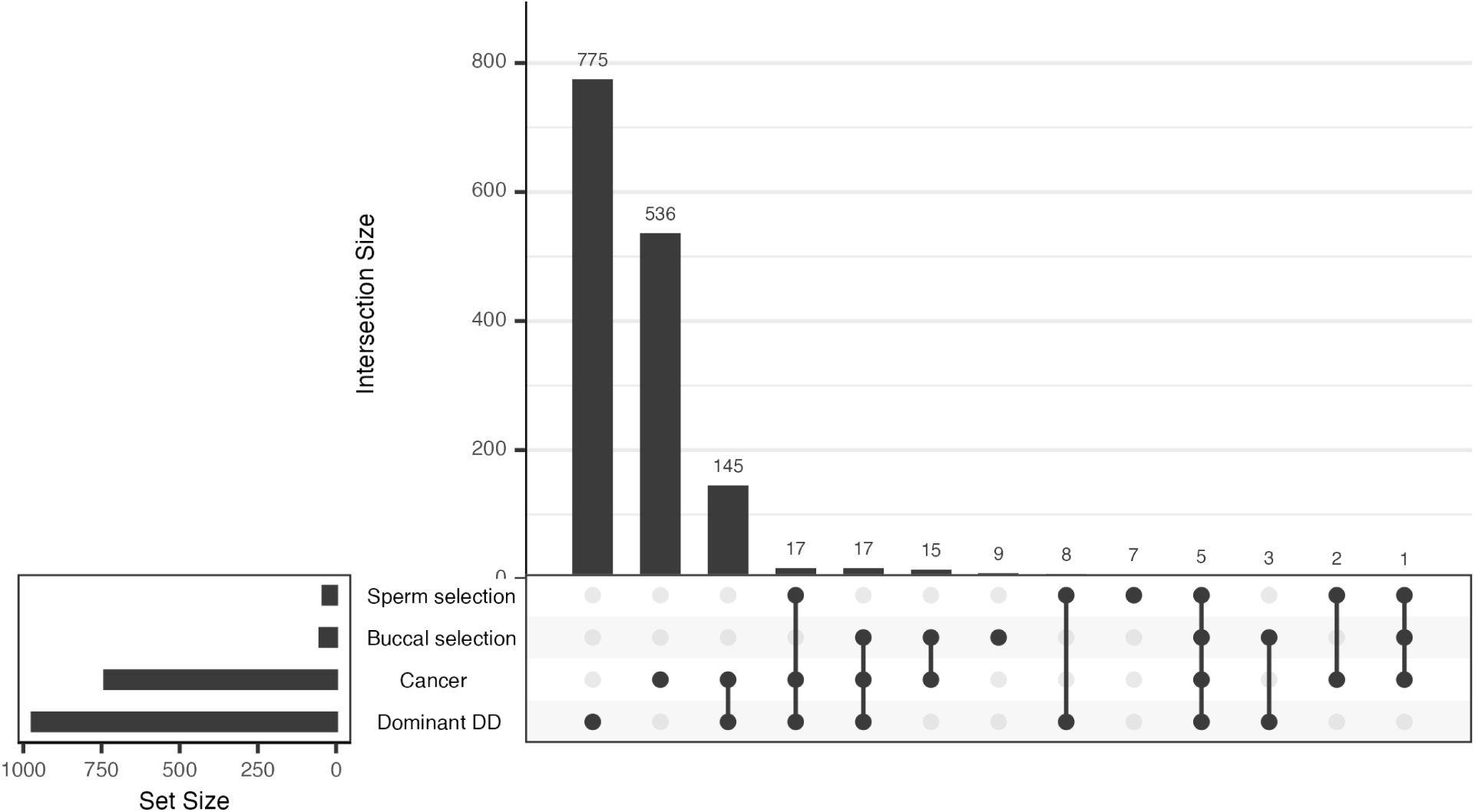
Upset plot showing overlap of dominant DD genes with genes showing evidence of somatic selection in cancer, sperm, and buccal tissue.

**Figure S2:**
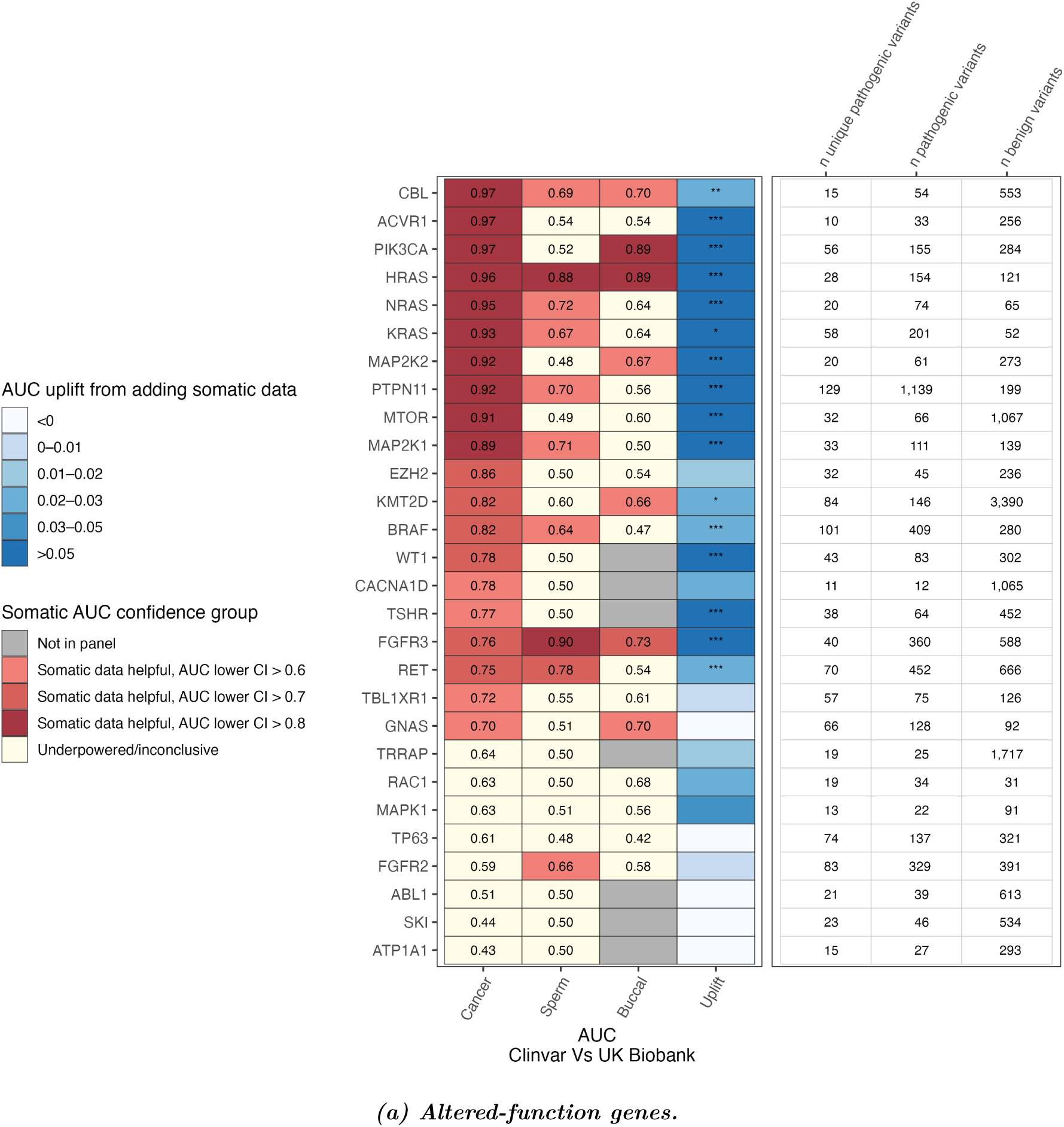
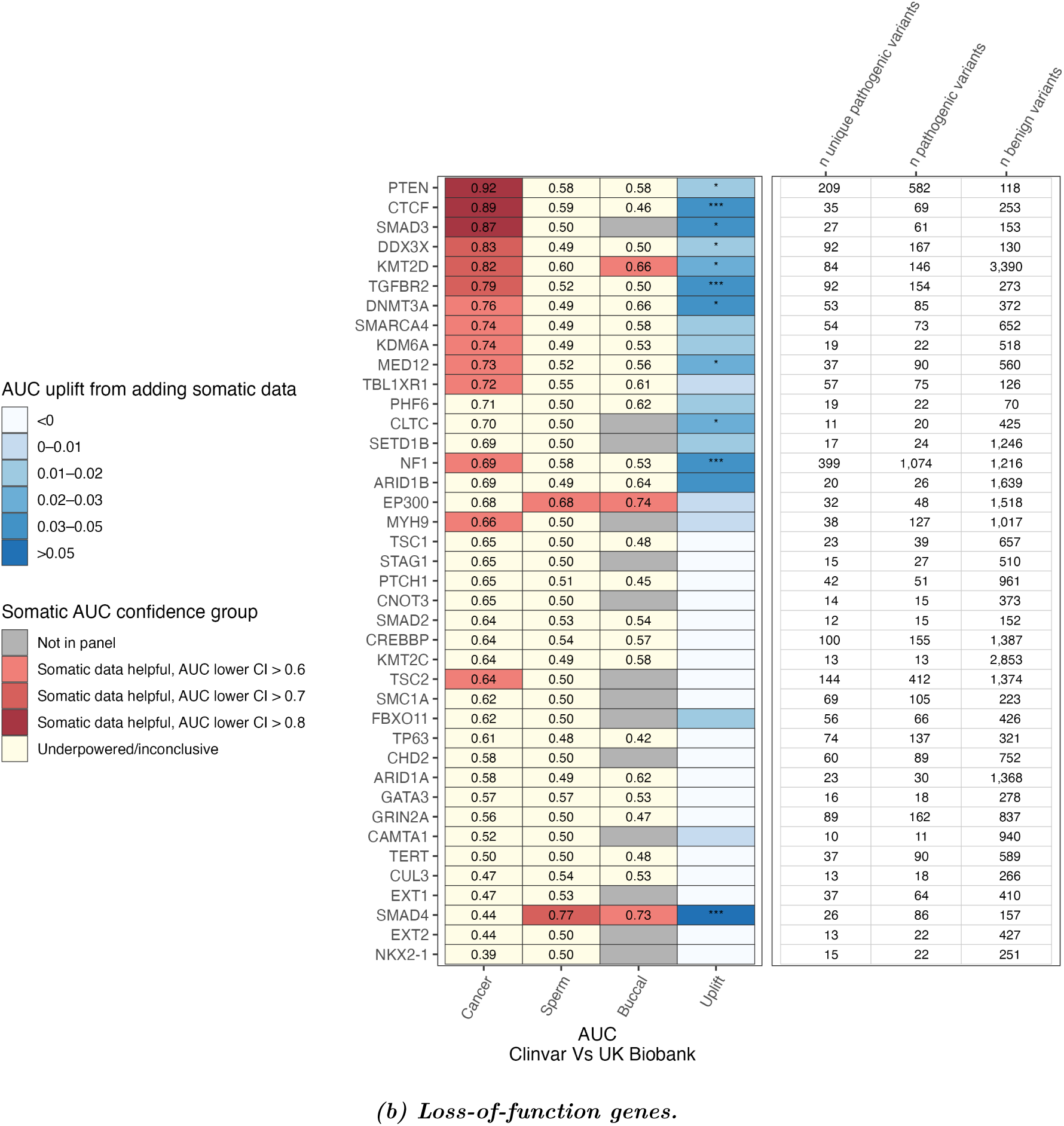
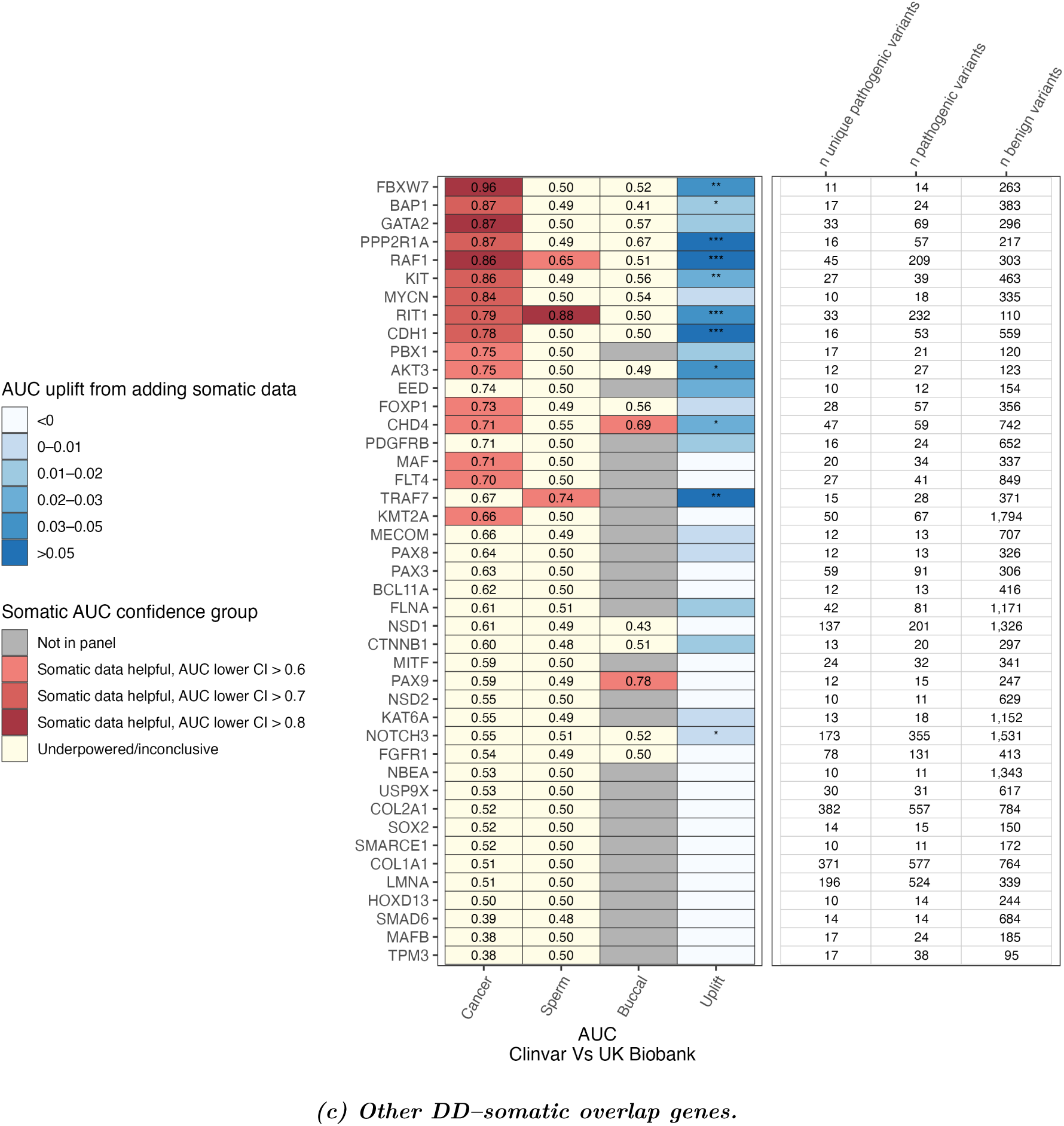
Heterogeneity in performance of different somatic data sources across mechanism-defined gene groups. (a) Altered-function genes. (b) Loss-of-function genes. (c) Other dominant DD genes with evidence of somatic selection.

**Figure S3:**
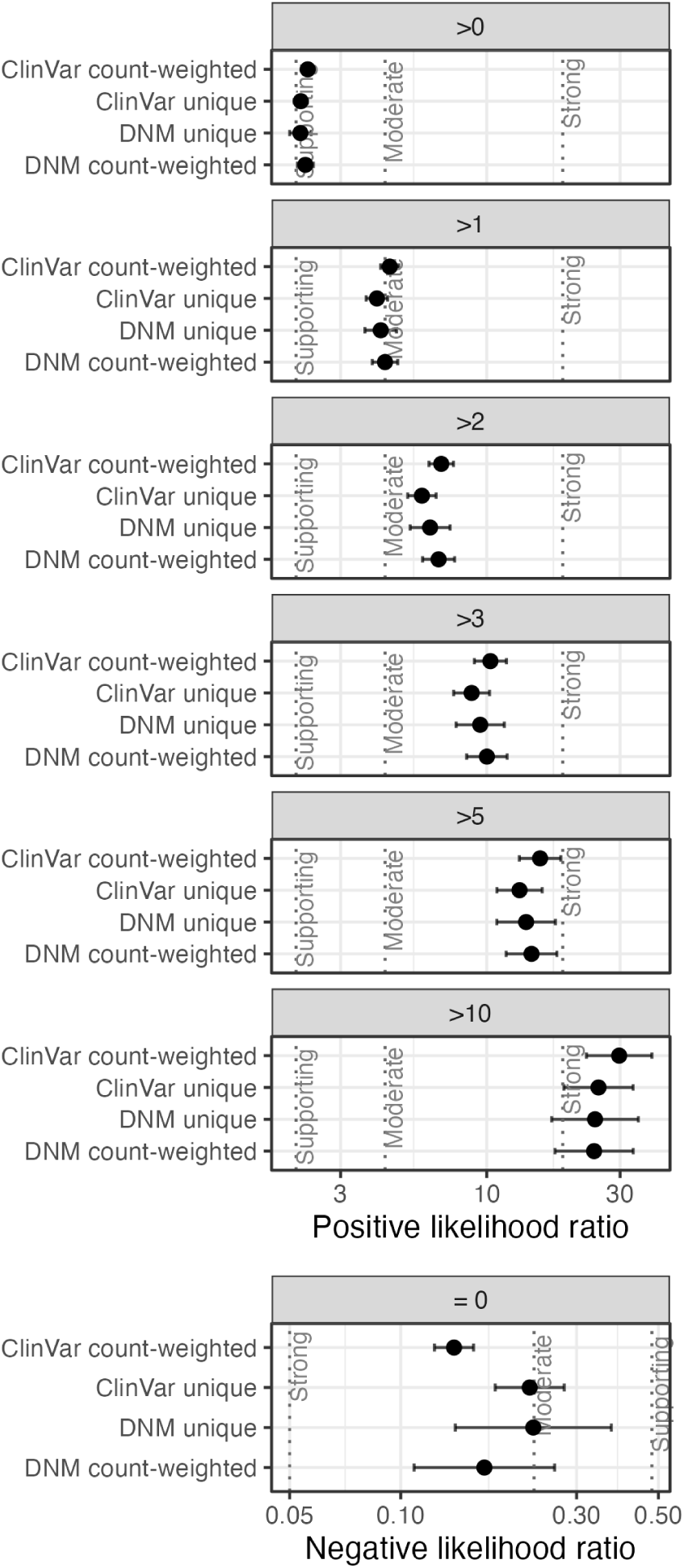
COSMIC missense codon count likelihood-ratio thresholds stratified by pathogenic variant data type.

## Notes

### Competing Interest Statement

The authors have declared no competing interest.

## References

[1] Melissa J Landrum, Jennifer M Lee, Mark Benson, Garth R Brown, Chen Chao, Shanmuga Chitipiralla, Bihan Gu, Joshua Hart, David Hoffman, Woobin Jang, et al. Clinvar: improving access to variant interpretations and supporting evidence. Nucleic Acids Research, 46(D1):D1062–D1067, January 2018. doi: 10.1093/nar/gkx1153.

[2] Lukas Gerasimavicius, Benjamin J Livesey, and Joseph A Marsh. Loss-of-function, gain-of-function and dominant-negative mutations have profoundly different effects on protein structure. Nature Communications, 13(1):3895, July 2022.

[3] Jasmin J Hopkins, Matthew N Wakeling, Matthew B Johnson, Sarah E Flanagan, and Thomas W Laver. REVEL is better at predicting pathogenicity of loss-of-function than gain-of-function variants. Human Mutation, 2023(1):1–6, December 2023.

[4] Suzanne Schubbert, Kevin Shannon, and Gideon Bollag. Hyperactive ras in developmental disorders and cancer. Nature Reviews Cancer, 7(4):295–308, April 2007.

[5] Bengi Ruken Yavuz, M Kaan Arici, Habibe Cansu Demirel, Chung-Jung Tsai, Hyunbum Jang, Ruth Nussinov, and Nurcan Tuncbag. Neurodevelopmental disorders and cancer networks share pathways, but differ in mechanisms, signaling strength, and outcome. NPJ Genom. Med., 8(1):37, November 2023.

[6] Bushra Haque, David Cheerie, Amy Pan, Meredith Curtis, Thomas Nalpathamkalam, Jimmy Nguyen, Celine Salhab, Bhooma Thiruvahindrapuram, Jade Zhang, Madeline Couse, Taila Hartley, Michelle M Morrow, E Magda Price, Susan Walker, David Malkin, Frederick P Roth, and Gregory Costain. Leveraging cancer mutation data to inform the pathogenicity classification of germline missense variants. PLoS Genetics, 21(1):e1011540, January 2025. doi: 10.1371/journal.pgen.1011540.

[7] Sean V Tavtigian, Marc S Greenblatt, Steven M Harrison, Robert L Nussbaum, Sankaranarayanan A Prabhu, Karen M Boucher, and Leslie G Biesecker. Modeling the ACMG/AMP variant classification guidelines as a bayesian classification framework. Genetics in Medicine, 20(9):1054–1060, 2018. doi: 10.1038/gim.2017.210.

[8] Michael F Walsh, Deborah I Ritter, Chimene Kesserwan, Dmitriy Sonkin, Debyani Chakravarty, Elizabeth Chao, Rajarshi Ghosh, Yelena Kemel, Gang Wu, Kristy Lee, Shashikant Kulkarni, Dale Hedges, Diana Mandelker, Ozge Ceyhan-Birsoy, Minjie Luo, Michael Drazer, Liying Zhang, Kenneth Offit, and Sharon E Plon. Integrating somatic variant data and biomarkers for germline variant classification in cancer predisposition genes. Human Mutation, 39(11):1542–1552, November 2018. doi: 10.1002/humu.23640.

[9] Matthew T Chang, Saurabh Asthana, Sizhi Paul Gao, Byron H Lee, Jocelyn S Chapman, Cyriac Kandoth, Jianjiong Gao, Nicholas D Socci, David B Solit, Adam B Olshen, Nikolaus Schultz, and Barry S Taylor. Identifying recurrent mutations in cancer reveals widespread lineage diversity and mutational specificity. Nature Biotechnology, 34(2):155–163, 2016. doi: 10.1038/nbt.3391.URL https://www.cancerhotspots.org. CancerHotspots.org (accessed 2026-03-13).

[10] Clinical Genome Resource (ClinGen) RASopathy Variant Curation Expert Panel. Clingen rasopathy expert panel specifications to the ACMG/AMP variant interpretation guidelines (version 1) and gene list. Criteria Specification Registry (CSpec), 2021. URL https://genboree.org/cspec/ui/svi/svi/GN004. Accessed 2026-03-13.

[11] Inigo Martincorena, Azim Roshan, Moritz Gerstung, Peter Ellis, Peter Van Loo, Stuart McLaren, David C Wedge, Anthony Fullam, Ludmil B Alexandrov, Jose M C Tubio, et al. High burden and pervasive positive selection of somatic mutations in normal human skin. Science, 348(6237):880–886, May 2015. doi: 10.1126/science.aaa6806.

[12] Inigo Martincorena, Joanna C Fowler, Agnieszka Wabik, Andrew R J Lawson, Federico Abascal, Michael W J Hall, Alex Cagan, Kazuhiro Murai, Krishnaa Mahbubani, Michael R Stratton, et al. Somatic mutant clones colonize the human esophagus with age. Science, 362(6417):911–917, October 2018. doi: 10.1126/science.aau3879.

[13] Andrew R J Lawson, Federico Abascal, Pantelis A Nicola, Stefanie V Lensing, Amy L Roberts, Georgios Kalantzis, Adrian Baez-Ortega, Natalia Brzozowska, Julia S El-Sayed Moustafa, Dovile Vaitkute, Belma Jakupovic, Ayrun Nessa, Samuel Wadge, Marc F Osterdahl, Anna L Paterson, et al. Somatic mutation and selection at population scale. Nature, 647(8089):411–420, 2025. doi: 10.1038/s41586-025-09584-w.

[14] Matthew D C Neville, Andrew R J Lawson, Rashesh Sanghvi, Federico Abascal, My H Pham, Alex Cagan, Pantelis A Nicola, Tetyana Bayzetinova, Adrian Baez-Ortega, Kirsty Roberts, Marianne B Jakobsen, Parvaz Fasihi, et al. Sperm sequencing reveals extensive positive selection in the male germline. Nature, 647(8089):421–428, 2025. doi: 10.1038/s41586-025-09448-3.

[15] Federico Abascal, Luke M R Harvey, Emily Mitchell, Andrew R J Lawson, Stefanie V Lensing, Peter Ellis, Andrew J C Russell, Raul E Alcantara, Adrian Baez-Ortega, Yichen Wang, Eugene Jing Kwa, Henry Lee-Six, Alex Cagan, Tim H H Coorens, Michael Spencer Chapman, Sigurgeir Olafsson, Steven Leonard, David Jones, Heather E Machado, Megan Davies, Nina F Øbro, Krishnaa T Mahubani, Kieren Allinson, Moritz Gerstung, Kourosh Saeb-Parsy, David G Kent, Elisa Laurenti, Michael R Stratton, Raheleh Rahbari, Peter J Campbell, Robert J Osborne, and Iñigo Martincorena. Somatic mutation landscapes at single-molecule resolution. Nature, 593(7859):405–410, 2021.

[16] EMBL-EBI. Gene2phenotype (g2p). Website. URL https://www.ebi.ac.uk/gene2phenotype/. Accessed 2025-04-16.

[17] T. Michael Yates, Morad Ansari, Louise Thompson, Sarah E. Hunt, Elena Cibrian Uhalte, Rachel J. Hobson, Joseph A. Marsh, Caroline F. Wright, Helen V. Firth, et al. Curating genomic diseasegene relationships with Gene2Phenotype (G2P). Genome Medicine, 16:127, November 2024. doi:10.1186/s13073-024-01398-1.

[18] Zbyslaw Sondka, Samuel Bamford, Charlotte G Cole, Sari A Ward, Ian Dunham, and Simon A Forbes. The cosmic cancer gene census: describing genetic dysfunction across all human cancers. Nature Reviews Cancer, 18(11):696–705, 2018. doi: 10.1038/s41568-018-0060-1.

[19] WikiPathways. Wikipathways pathway wp3844 (pi3k–mtor signaling pathway). Webpage (ComPath mirror). URL https://compath.scai.fraunhofer.de/pathway/wikipathways/WP3844. Accessed 2026-05-07.

[20] Kanehisa Laboratories. Kegg pathway: Mapk signaling pathway (hsa04010). Webpage. URL https://www.kegg.jp/pathway/hsa04010. Accessed 2026-03-23.

[21] Joanna Kaplanis, Kaitlin E Samocha, Luan H Wiel, Zhuozhi Zhang, Kristina J Arvai, Ryan Y Eberhardt, Giovanni Gallone, Sander H Lelieveld, Hilary C Martin, Jeremy F McRae, et al. Integrating healthcare and research genetic data empowers the discovery of 28 novel developmental disorders. Nature, 586 (7831):757–762, October 2020. doi: 10.1038/s41586-020-2832-5.

[22] Petr Danecek and Shane A McCarthy. BCFtools/csq: haplotype-aware variant consequences. Bioinformatics, 33(13):2037–2039, July 2017. Accessed 2026-04-10.

[23] Joseph D Szustakowski, Suganthi Balasubramanian, Erika Kvikstad, Shareef Khalid, Paola G Bronson, Ariella Sasson, Emily Wong, Daren Liu, J Wade Davis, Carolina Haefliger, A Katrina Loomis, Rajesh Mikkilineni, Hyun Ji Noh, Samir Wadhawan, Xiaodong Bai, Alicia Hawes, Olga Krasheninina, Ricardo Ulloa, Alex E Lopez, Erin N Smith, Jeffrey F Waring, Christopher D Whelan, Ellen A Tsai, John D Overton, William J Salerno, Howard Jacob, Sandor Szalma, Heiko Runz, Gregory Hinkle, Paul Nioi, Slavé Petrovski, Melissa R Miller, Aris Baras, Lyndon J Mitnaul, Jeffrey G Reid, and UKB-ESC Research Team. Advancing human genetics research and drug discovery through exome sequencing of the UK biobank. Nature Genetics, 53(7):942–948, July 2021. Published online 2021-07-28 (accessed 2026-05-06).

[24] Eugene J Gardner, Katherine A Kentistou, Stasa Stankovic, Samuel Lockhart, Eleanor Wheeler, Felix R Day, Nicola D Kerrison, Nicholas J Wareham, Claudia Langenberg, Stephen O’Rahilly, Ken K Ong, and John R B Perry. Damaging missense variants in IGF1R implicate a role for IGF-1 resistance in the etiology of type 2 diabetes. Cell Genomics, 2(12), December 2022. Published online 2022-12-14 (accessed 2026-05-08).

[25] Martin Kircher, Daniela M Witten, Preti Jain, Brian J O’Roak, Gregory M Cooper, and Jay Shendure. A general framework for estimating the relative pathogenicity of human genetic variants. Nature Genetics, 46(3):310–315, 2014. doi: 10.1038/ng.2892.

[26] Nicholas M Ioannidis, Joseph H Rothstein, Vikas Pejaver, Shashikant Middha, Shannon K McDonnell, Sujata Baheti, Alicia Musolf, Qing Li, Emily Holzinger, David Karyadi, et al. Revel: An ensemble method for predicting the pathogenicity of rare missense variants. American Journal of Human Genetics, 99(4):877–885, September 2016. doi: 10.1016/j.ajhg.2016.08.016.

[27] Rose Orenbuch, Courtney A Shearer, Aaron W Kollasch, Aviv D Spinner, Thomas Hopf, Lood van Niekerk, Dinko Franceschi, Mafalda Dias, Jonathan Frazer, and Debora S Marks. Proteome-wide model for human disease genetics. Nature Genetics, 57(12):3165–3174, December 2025. Published online 2025-12-24 (accessed 2026-06-16).

[28] Jun Cheng, Guido Novati, Joshua Pan, Clare Bycroft, Akvile Žemgulyte, Taylor Applebaum, Alexander Pritzel, Lai Hong Wong, Michal Zielinski, Tobias Sargeant, Rosalia G Schneider, Andrew W Senior, John Jumper, Demis Hassabis, Pushmeet Kohli, and Žiga Avsec. Accurate proteome-wide missense variant effect prediction with AlphaMissense. Science, 381(6664):eadg7492, 2023. doi: 10.1126/science.adg7492.

[29] Nadav Brandes, Grant Goldman, Charlotte H Wang, Chun Jimmie Ye, and Vasilis Ntranos. Genome-wide prediction of disease variant effects with a deep protein language model. Nature Genetics, 55(9): 1512–1522, 2023. doi: 10.1038/s41588-023-01465-0.

[30] Laksshman Sundaram, Hong Gao, Samskruthi Reddy Padigepati, Jeremy F McRae, Yanjun Li, Jack A Kosmicki, Nondas Fritzilas, Jörg Hakenberg, Anindita Dutta, John Shon, Jinbo Xu, Serafim Batzoglou, Xiaolin Li, and Kyle Kai-How Farh. Predicting the clinical impact of human mutation with deep neural networks. Nature Genetics, 50(8):1161–1170, 2018. doi: 10.1038/s41588-018-0167-z.

[31] Edison Scientific. Edison scientific platform. Website. URL https://platform.edisonscientific.com/. Accessed 2026-04-10.

[32] Alan F Rubin, Jeremy Stone, Aisha Haley Bianchi, Benjamin J Capodanno, Estelle Y Da, Mafalda Dias, Daniel Esposito, Jonathan Frazer, Yunfan Fu, Sally B Grindstaff, Matthew R Harrington, Iris Li, Abbye E McEwen, Joseph K Min, Nick Moore, Olivia G Moscatelli, Jesslyn Ong, Polina V Polunina, Joshua E Rollins, Nathan J Rollins, Ashley E Snyder, Amy Tam, Matthew J Wakefield, Shenyi Sunny Ye, Lea M Starita, Vanessa L Bryant, Debora S Marks, and Douglas M Fowler. MaveDB 2024: a curated community database with over seven million variant effects from multiplexed functional assays. Genome Biology, 26(1):13, January 2025. doi: 10.1186/s13059-025-03476-y.

[33] Elizabeth J Radford, Hong-Kee Tan, Malin H L Andersson, James D Stephenson, Eugene J Gardner, Holly Ironfield, Andrew J Waters, Daniel Gitterman, Sarah Lindsay, Federico Abascal, Iñigo Martincorena, Anna Kolesnik-Taylor, Elise Ng-Cordell, Helen V Firth, Kate Baker, John R B Perry, David J Adams, Sebastian S Gerety, and Matthew E Hurles. Saturation genome editing of DDX3X clarifies pathogenicity of germline and somatic variation. Nature Communications, 14(1):7702, December 2023. doi: 10.1038/s41467-023-43041-4.

[34] Ziyuan Jiang, Anne E van Vlimmeren, Deepti Karandur, Alyssa Semmelman, and Neel H Shah. Deep mutational scanning of the multi-domain phosphatase SHP2 reveals mechanisms of regulation and pathogenicity. Nat. Commun., 16(1):5464, 2025.

[35] Timothy L. Mighell, Sarah Evans-Dutson, and Brian J. O’Roak. A saturation mutagenesis approach to understanding PTEN lipid phosphatase activity and genotype-phenotype relationships. American Journal of Human Genetics, 102(5):943–955, May 2018. doi: 10.1016/j.ajhg.2018.03.018.

[36] Kenneth A. Matreyek, Lea M. Starita, Jeffrey J. Stephany, Blake Martin, Michael A. Chiasson, Victoria E. Gray, Martin Kircher, Akaki Khechaduri, Jennifer N. Dines, Ronald J. Hause, et al. Multiplex assessment of protein variant abundance by massively parallel sequencing. Nature Genetics, 50:874–882, 2018. doi: 10.1038/s41588-018-0122-z.

[37] Kenneth A Matreyek, Jason J Stephany, Ethan Ahler, and Douglas M Fowler. Integrating thousands of PTEN variant activity and abundance measurements reveals variant subgroups and new dominant negatives in cancers. 13:165.

[38] Frank Hidalgo, Laura M Nocka, Neel H Shah, Kent Gorday, Naomi R Latorraca, Pradeep Bandaru, Sage Templeton, David Lee, Deepti Karandur, Jeffrey G Pelton, Susan Marqusee, David Wemmer, and John Kuriyan. A saturation-mutagenesis analysis of the interplay between stability and activation in ras. eLife, 11:e76595, March 2022. Article e76595.

[39] S. Chen, L. C. Francioli, J. K. Goodrich, R. L. Collins, M. Kanai, Q. Wang, J. Alföldi, N. A. Watts, C. Vittal, L. D. Gauthier, T. Poterba, M. W. Wilson, Y. Tarasova, W. Phu, R. Grant, M. T. Yohannes, Z. Koenig, Y. Farjoun, E. Banks, S. Donnelly, S. Gabriel, N. Gupta, S. Ferriera, C. Tolonen, S. Novod, L. Bergelson, D. Roazen, V. Ruano-Rubio, M. Covarrubias, C. Llanwarne, N. Petrillo, G. Wade, T. Jeandet, R. Munshi, K. Tibbetts, Genome Aggregation Database (gnomAD) Consortium, A. O’Donnell-Luria, M. Solomonson, C. Seed, A. R. Martin, M. E. Talkowski, H. L. Rehm, M. J. Daly, G. Tiao, B. M. Neale, D. G. MacArthur, and K. J. Karczewski. A genomic mutational constraint map using variation in 76,156 human genomes. Nature, 625:92–100, 2024. doi: 10.1038/s41586-023-06045-0.

[40] Hongjian Qi, Chengliang Dong, Wendy K Chung, Kai Wang, and Yufeng Shen. Deep genetic connection between cancer and developmental disorders. Hum. Mutat., 37(10):1042–1050, October 2016.

[41] Cathie Sudlow, John Gallacher, Naomi Allen, Valerie Beral, Paul Burton, John Danesh, Paul Downey, Paul Elliott, Jane Green, Martin Landray, et al. Uk biobank: an open access resource for identifying the causes of a wide range of complex diseases of middle and old age. PLoS Medicine, 12(3):e1001779, March 2015. doi: 10.1371/journal.pmed.1001779.

[42] Katherine A Wood, R Spencer Tong, Marialetizia Motta, Viviana Cordeddu, Eleanor R Scimone, Stephen J Bush, Dale W Maxwell, Eleni Giannoulatou, Viviana Caputo, Alice Traversa, Cecilia Mancini, Giovanni B Ferrero, Francesco Benedicenti, Paola Grammatico, Daniela Melis, Katharina Steindl, Nicola Brunetti-Pierri, Eva Trevisson, Andrew Om Wilkie, Angela E Lin, Valerie Cormier-Daire, Stephen Rf Twigg, Marco Tartaglia, and Anne Goriely. SMAD4 mutations causing myhre syndrome are under positive selection in the male germline. Am. J. Hum. Genet., 111(9):1953–1969, September 2024.

[43] K. J. Champion, C. Bunag, A. L. Estep, J. R. Jones, C. H. Bolt, R. C. Rogers, K. A. Rauen, and D. B. Everman. Germline mutation in BRAF codon 600 is compatible with human development: de novo p.V600G mutation identified in a patient with CFC syndrome. Clinical Genetics, 79(5):468–474, May 2011.

[44] Cigdem Sevim Bayrak, David Stein, Aayushee Jain, Kumardeep Chaudhary, Girish N Nadkarni, Tielman T Van Vleck, Anne Puel, Stephanie Boisson-Dupuis, Satoshi Okada, Peter D Stenson, David N Cooper, Avner Schlessinger, and Yuval Itan. Identification of discriminative gene-level and protein-level features associated with pathogenic gain-of-function and loss-of-function variants. American Journal of Human Genetics, 108(12):2301–2318, December 2021.

[45] Harriet Copeland, Karen J Low, Sarah L Wynn, Ayesha Ahmed, Victoria Arthur, Meena Balasubramanian, Katya Bennett, Jonathan Berg, Marta Bertoli, Lisa Bryson, Catrin Bucknall, Jamie Campbell, Kate Chandler, Jaynee Chauhan, Amy Clarkson, Rachel Coles, Hector Conti, Philandra Costello, Tessa Coupar, Amy Craig, John Dean, Amy Dillon, Abhijit Dixit, Kathryn Drew, Jacqueline Eason, Francesca Forzano, Nicola Foulds, Alice Gardham, Neeti Ghali, Andrew Green, William Hanna, Rachel Harrison, Mairead Hegarty, Jenny Higgs, Muriel Holder, Rachel Irving, Vani Jain, Katie Johnson, Rachel Jolley, Wendy D Jones, Gabriela Jones, Shelagh Joss, Ruta Kalinauskiene, Farah Kanani, Karl Kavanagh, Mahmudur Khan, Naz Khan, Emma Kivuva, Nayana Lahiri, Neeta Lakhani, Anne Lampe, Sally Ann Lynch, Sahar Mansour, Alice Marsden, Hannah Massey, Shane McKee, Shehla Mohammed, Swati Naik, Mithushanaa Nesarajah, Ruth Newbury-Ecob, Fiona Osborne, Michael J Parker, Jenny Patterson, Caroline Pottinger, Matina Prapa, Katrina Prescott, Shauna Quinn, Jessica A Radley, Sarah Robart, Alison Ross, Giulia Rosti, Francis H Sansbury, Ajoy Sarkar, Claire Searle, Nora Shannon, Debbie Shears, Sarah Smithson, Helen Stewart, Mohnish Suri, Shereen Tadros, Rachel Theobald, Rhian Thomas, Olga Tsoulaki, Pradeep Vasudevan, Maribel Verdesoto Rodriguez, Emma Vittery, Sinead Whyte, Emily Woods, Thomas Wright, David Zocche, Helen V Firth, Caroline F Wright, and DDD Study. Large-scale evaluation of outcomes after a genetic diagnosis in children with severe developmental disorders. Genet. Med. Open, 2:101864, October 2024.

[46] Sue Richards, Nazneen Aziz, Sherri Bale, David Bick, Soma Das, Julie Gastier-Foster, Wayne W Grody, Madhuri Hegde, Elaine Lyon, Elaine Spector, Karl Voelkerding, Heidi L Rehm, and ACMG Laboratory Quality Assurance Committee. Standards and guidelines for the interpretation of sequence variants: a joint consensus recommendation of the american college of medical genetics and genomics and the association for molecular pathology. Genetics in Medicine, 17(5):405–424, May 2015.

[47] Vikas Pejaver, Alicia B Byrne, Bing-Jian Feng, Kymberleigh A Pagel, Sean D Mooney, Rachel Karchin, Anne O’Donnell-Luria, Steven M Harrison, Sean V Tavtigian, Marc S Greenblatt, Leslie G Biesecker, Predrag Radivojac, Steven E Brenner, and ClinGen Sequence Variant Interpretation Working Group. Calibration of computational tools for missense variant pathogenicity classification and ClinGen recommendations for PP3/BP4 criteria. American Journal of Human Genetics, 109(12):2163–2177, December 2022.

[48] Timothy Bergquist, Sarah L Stenton, Emily A W Nadeau, Alicia B Byrne, Marc S Greenblatt, Steven M Harrison, Sean V Tavtigian, Anne O’Donnell-Luria, Leslie G Biesecker, Predrag Radivojac, Steven E Brenner, Vikas Pejaver, and ClinGen Sequence Variant Interpretation Working Group. Calibration of additional computational tools expands ClinGen recommendation options for variant classification with PP3/BP4 criteria. Genetics in Medicine, 27(6):101402, June 2025.

[49] Julia Foreman, Daniel Perrett, Erica Mazaika, Sarah E Hunt, James S Ware, and Helen V Firth. DECIPHER: Improving genetic diagnosis through dynamic integration of genomic and clinical data. Annual Review of Genomics and Human Genetics, 24:151–176, August 2023. Published online 2023-08-25 (accessed 2026-06-16).

